# A bistable inhibitory OptoGPCR for multiplexed optogenetic control of neural circuits

**DOI:** 10.1101/2023.07.01.547328

**Authors:** Jonas Wietek, Adrianna Nozownik, Mauro Pulin, Inbar Saraf-Sinik, Noa Matosevich, Daniela Malan, Bobbie J. Brown, Julien Dine, Rivka Levy, Anna Litvin, Noa Regev, Suraj Subramaniam, Eyal Bitton, Asaf Benjamin, Bryan A. Copits, Philipp Sasse, Benjamin R. Rost, Dietmar Schmitz, Peter Soba, Yuval Nir, J. Simon Wiegert, Ofer Yizhar

**Affiliations:** Department of Brain Sciences, Weizmann Institute of Science, Rehovot, Israel; Department of Molecular Neuroscience, Weizmann Institute of Science, Rehovot, Israel; Center for Molecular Neurobiology, Hamburg, Germany; Present address: Paris Brain Institute, Institut du Cerveau (ICM), CNRS UMR 7225, INSERM U1127, Sorbonne Université, Paris, France; Present address: Laboratory of Sensory Processing, Brain Mind Institute, Faculty of Life Sciences, Ecole Polytechnique Fédérale de Lausanne (EPFL), Lausanne, Switzerland; Sagol school of neuroscience, Tel Aviv University, Tel Aviv, Israel; Institut für Physiologie I, Universität Bonn, Bonn, Germany; Washington University Pain Center, Department of Anesthesiology, Washington University School of Medicine, St. Louis, MO, USA; Present address: Boehringer Ingelheim Pharma GmbH & Co. KG; CNS Diseases, Biberach an der Riss, Germany; Department of Physiology and Pharmacology, Tel Aviv University, Tel Aviv, Israel; German Center for Neurodegenerative Diseases (DZNE), Berlin, Germany; Neuroscience Research Center, Charité – Universitätsmedizin Berlin, Berlin, Germany; Bernstein Center for Computational Neuroscience, Berlin, Germany; Einstein Center for Neurosciences Berlin, Berlin, Germany; Max-Delbrück Center for Molecular Medicine, Berlin, Germany; Institute of Physiology and Pathophysiology, Friedrich-Alexander-Universität Erlangen-Nürnberg, Erlangen, Germany; LIMES-Institute, University of Bonn, Bonn, Germany; Department of Biomedical Engineering, Faculty of Engineering, Tel Aviv University, Tel Aviv, Israel; Present address: MCTN, Medical Faculty Mannheim of the University of Heidelberg, Mannheim, Germany

## Abstract

Information is transmitted between brain regions through the release of neurotransmitters from long-range projecting axons. Understanding how the activity of such long-range connections contributes to behavior requires efficient methods for reversibly manipulating their function. Chemogenetic and optogenetic tools, acting through endogenous G-protein coupled receptor (GPCRs) pathways, can be used to modulate synaptic transmission, but existing tools are limited in sensitivity, spatiotemporal precision, or spectral multiplexing capabilities. Here we systematically evaluated multiple bistable opsins for optogenetic applications and found that the *Platynereis dumerilii* ciliary opsin (*Pd*CO) is an efficient, versatile, light-activated bistable GPCR that can suppress synaptic transmission in mammalian neurons with high temporal precision *in-vivo*. *Pd*CO has superior biophysical properties that enable spectral multiplexing with other optogenetic actuators and reporters. We demonstrate that *Pd*CO can be used to conduct reversible loss-of-function experiments in long-range projections of behaving animals, thereby enabling detailed synapse-specific functional circuit mapping.

## Main

Even the simplest behaviors are coordinated by neural activity spanning multiple brain regions. Long-range projecting axons facilitate this by forming synaptic connections in distant circuits in the brain and periphery. Understanding the roles of these long-range connections in circuit activity and behavior requires techniques that allow selective manipulation of their function. While optogenetic tools allow the manipulation of neural firing with high temporal and spatial precision^1^, projection neurons often target several downstream regions via highly branched axonal collaterals^2,3^. Thus, manipulating the neuronal soma may result in a partial or even misleading picture of their contribution to circuit function. Instead, directly targeting the synaptic terminals of long-range connections can provide fine-grained insight into the role of specific neuronal pathways. However, direct suppression of synaptic terminal function poses unique challenges. Inhibitory optogenetic tools, such as the microbial light-driven ion pumps, have traditionally been used to silence synaptic transmission at axonal terminals^4–8^. Their inhibitory effect on synaptic release, however, is not only partial and short-lived, it can also induce unintended paradoxical effects, such as an increase of spontaneous transmitter release^9^. Chloride-conducting channelrhodopsins also proved unsuitable for synaptic silencing, as they depolarize the axon and can trigger antidromic firing due to the high chloride reversal potential in this subcellular compartment^9–11^. In contrast, targeting inhibitory GPCR pathways with bistable rhodopsins was shown to be effective for attenuating synaptic release in a projection-specific manner^12–14^.

We and others have recently shown that exogenously expressed light-activated animal rhodopsins can transiently inhibit synaptic transmission, by coupling to endogenous inhibitory G-proteins. While visual rhodopsins can be expressed in neurons and used to suppress synaptic release^12^, these photoreceptors can undergo bleaching (i.e. they lose their light-sensitive chromophore retinal), which reduces their efficacy under sustained illumination^15^. In contrast, bleaching-resistant non-visual rhodopsins have recently gained attention as light-activated tools for suppression of presynaptic release^16^. Like endogenous inhibitory GPCRs, these light-activated GPCRs (optoGPCRs) trigger the opening of G protein-coupled inwardly rectifying potassium (GIRK) channels, activate the Gα_i/o_ signaling pathway (Fig. 1a) and efficiently suppress synaptic transmission through the inhibition of voltage-gated calcium channels (VGCCs) and the SNARE (soluble N-ethylmaleimide-sensitive-factor attachment receptor)-mediated fusion of synaptic vesicles with the presynaptic membrane^17,18^. Activated Gα_i_ subunits can also reduce cyclic adenosine monophosphate (cAMP) production by adenylate cyclases (ACs), indirectly decreasing cAMP-dependent neurotransmission^19–21^.

**Figure 1:**
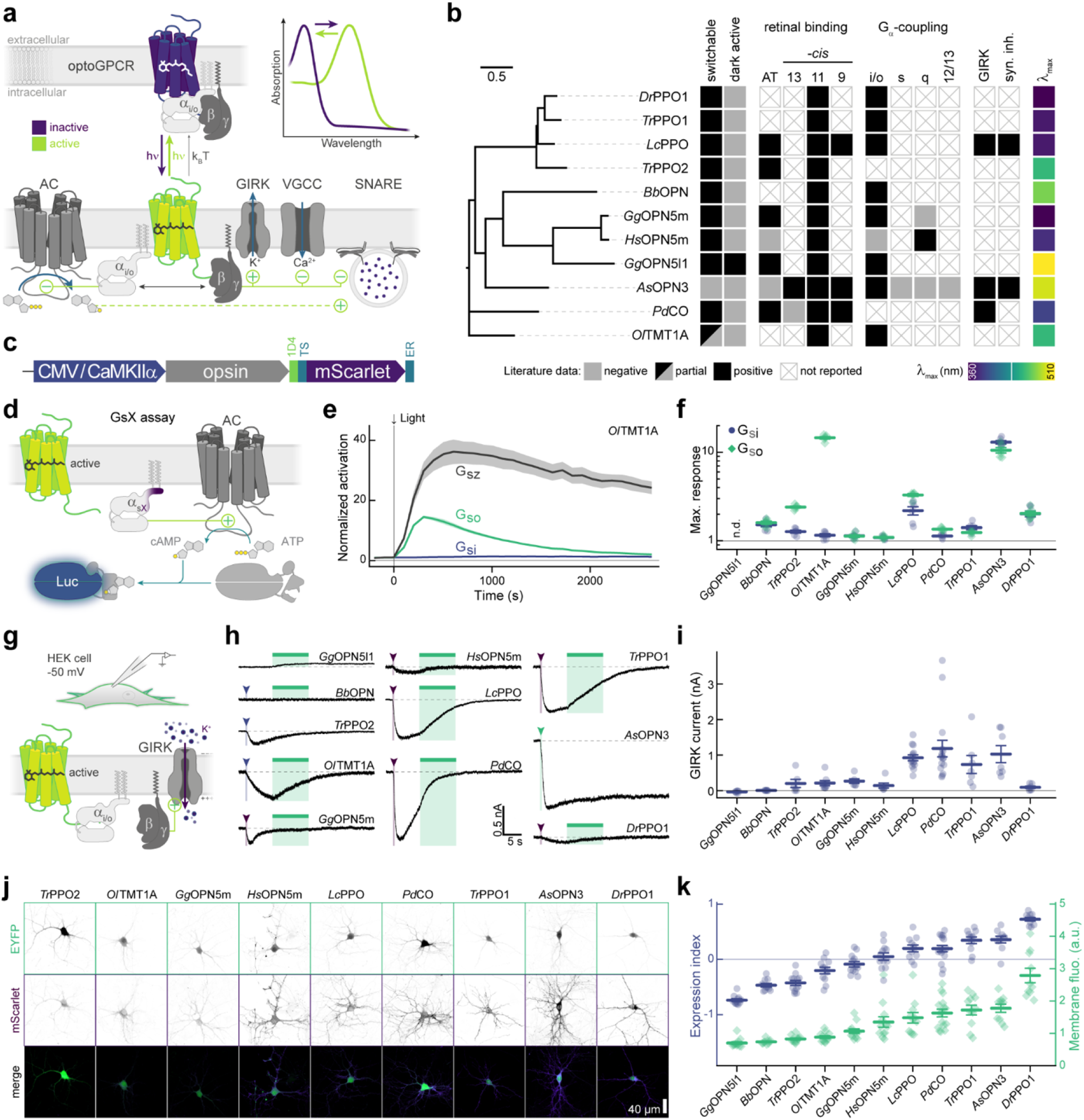
Benchmarking of inhibitory optoGPCR candidates. **a**, Scheme of inhibitory optoGPCRs that couple via the G_i/o_-pathway. A dark, inactive optoGPCR bound to the heterotrimeric Gαβγ protein is shown. Once the optoGPCR is activated by light, the heterotrimeric G-protein separates into the active Gα and Gβγ subunits. Gβγ activates G-protein coupled inward rectifying potassium channels (GIRK), inhibits voltage-gated calcium channels (VGCCs) and may interfere with the SNARE (soluble *N*-ethylmaleimide-sensitive-factor attachment receptor) vesicle fusion apparatus. The Gα subunit inhibits adenylyl cyclases (ACs), thus reducing production of cAMP. OptoGPCRs can relax thermally (k_B_T) to the non-signaling ground state. If their active state is spectrally separated from the ground state (see inset: absorption), absorption of a second photon with longer wavelength (hν) can terminate the signaling activity. **b**, Overview of optoGPCR candidates investigated in this study. Left, phylogenetic tree of optoGPCRs. Right, properties of optoGPCRs as reported in available literature. AT: all-*trans*. **c**, Design of DNA constructs used for initial characterization. **d**, Characterization of each optoGPCR’s Gα-protein signaling profile using the GsX assay. Photoactivation of the optoGPCR activates a chimeric Gα_s_ subunit harboring the C-terminus from one other Gα-protein. This chimeric Gα-protein activates an adenylate cyclase (AC), which generates cAMP from ATP. cAMP activates a cAMP-dependent luciferase (Luc) whose bioluminescence is detected by a plate reader. **e**, Time course of averaged bioluminescence reads for Gα_sz_, Gα_si_, and Gα_so_ activation by *Ol*TMT1A. The bioluminescence signal was normalized to the signal of cells not expressing optoGPCR and to pre-illumination baseline. A 1 s 470 nm light pulse was used for activation. n= 6. **f**, Maximum bioluminescence response for light-activated optoGPCRs coupling to Gα_si_ (blue circles) and Gα_so_ (green diamonds). n.d.: not determined, n= 6. **g**, HEK-cell experiments to measure optoGPCR-evoked GIRK currents with whole-cell voltage-clamp recordings. **h**, Representative GIRK current traces recorded in HEK cells expressing the indicated optoGPCRs. Arrowheads and narrow bars indicate light application of 0.5 s, while wide bars indicate 10 s light activation (see Online Methods). **i**, Quantification of optoGPCR-evoked peak GIRK currents (n= 6 - 16). **j**, Maximum intensity projection images from confocal stacks of neurons expressing the nine best-expressing optoGPCRs fused to mScarlet together with a cytosolic EYFP. Inverted grayscale images are gamma corrected (1.25). The merged projections are false color-coded by green (EYFP) and duo intense purple (mScarlet) lookup tables. **k**, Quantification of membrane expression index (blue circles) and the membrane fluorescence (green diamonds) of each optoGPCR, determined from equatorial z-slices (n = 10 - 16). All data is shown as mean ± SEM.

Although progress has been made in developing inhibitory optoGPCRs, the existing tools are limited either in their spectral or temporal features. Two bistable rhodopsins, the trafficking-enhanced eOPN3 derived from *Anopheles stephensi*^13^ (referred to herewith as *As*OPN3) and the parapinopsin^14^ from the Japanese lamprey *Lethenteron camtschaticum (LcPPO)*, have recently been utilized as inhibitory optoGPCRs for presynaptic inhibition. The highly light-sensitive *As*OPN3 has a broad action spectrum that spans the entire UV-visible range^13^. However, *As*OPN3 activity cannot be rapidly reverted to the inactive (dark-adapted) state, and takes minutes to spontaneously recover to its non-signaling state^13,22^. *Lc*PPO can undergo photoswitching between its active and inactive states by different wavelengths, thus allowing better temporal control^23,24^. However, *Lc*PPO’s limitation lies in its UV maximum activation wavelength (∼370nm) and its broad inactivation spectrum^23,24^. These spectral properties restrict the wavelength range available for multiplexed applications with additional optogenetic actuators or fluorescence-based sensors. Especially for single-photon fiber photometry and miniature microscopy techniques, spectral multiplexing can be challenging with the current tools.

To improve and expand the capabilities of inhibitory optoGPCRs, we aimed for a new tool that retains the advantages of *As*OPN3 and *Lc*PPO but overcomes their limitations. We systematically screened a range of bistable opsins and evaluated their potential use as optoGPCRs based on their cellular biodistribution, spectral features and kinetic properties. Our screen revealed that the ciliary opsin 1 from *Platynereis dumerilii* (*Pd*CO)^25,26^ is a highly light-sensitive, bidirectionally switchable, versatile inhibitory optoGPCR. *Pd*CO expresses well in mammalian neurons and allows robust, high-efficiency, and rapidly switchable presynaptic silencing across various cell types and preparations. With its unique spectral features, including a red-shifted activation wavelength and a narrow inactivation spectrum, *Pd*CO is optimally suited for multiplexing with other optogenetic actuators and genetically-encoded sensors.

## Results

### Literature mining and functional benchmarking of optoGPCR candidates

We conducted a comprehensive literature search and identified a list of suitable optoGPCR candidates that could enable light-controlled inhibition of synaptic transmission. We collected information on retinal binding, spectral properties, switchability, G-protein coupling specificity, and activation of G-protein inward-rectifying potassium (GIRK) channels (Extended Data Fig. 1). Of the 32 described rhodopsins we selected for analysis, we identified 11 switchable variants that were most promising due to their coupling to Gα_i/o_ or activation of GIRK channels. Including the *As*OPN3 and *Lc*PPO for comparison (Fig. 1b), we conducted a three-part benchmark to characterize the functional properties of these optoGPCRs. Our goals were to: 1) profile their G-protein coupling specificity (Fig. 1d-f); 2) quantify GIRK channel currents induced by optoGPCRs activation (Fig. 1g-i); and 3) quantify their expression and membrane targeting in cultured neurons (Fig. 1j,k). Using the chimeric Gα bioluminescence assay (GsX^27^, Fig. 1d,e) we found that *Ol*TMT1A, *Lc*PPO, the PPOs from pufferfish (*Tr*PPO1/2), *Pd*CO, *As*OPN3, and *Dr*PPO1 couple to the inhibitory G_i/o_-protein family (G_i/o/t/z_, Fig. 1f). With the exception of *Tr*PPO2 that displayed additional coupling to the G_q/15_ and G_12/13_, and *Bb*OPN that showed non-selective G-protein coupling, we could not detect any G-protein activation other than for the G_i/o_ pathway (Extended Data Fig. 2).

Next, we expressed the optoGPCR candidates in human HEK293 cells that constitutively express GIRK2.1 channels, and measured the resulting G_βγ_-activated GIRK currents by whole-cell patch-clamp electrophysiology (Fig. 1g,h). *Lc*PPO, *Pd*CO, *Tr*PPO1, and *As*OPN3 produced the largest GIRK current amplitudes (>700 pA), while the other variants induced currents smaller than 270 pA (Fig. 1i). *Bb*OPN did not produce detectable light-induced currents, and *Gg*OPN5L1 displayed a small inhibition of GIRK currents, which is consistent with its reported dark-activity^28^. We also measured optoGPCR-evoked GIRK activation in cultured neurons, where all optoGPCR except *Bb*OPN and the OPN5 variants showed light-evoked GIRK conductance (Extended Data Fig. 4).

To quantify the expression level and membrane targeting of the optoGPCRs, we co-expressed each of the opsins with a cell-filling EYFP fluorophore and used confocal microscopy to measure protein expression and membrane targeting (Fig. 1j and Extended Data Fig. 3). In line with the electrophysiological recordings in HEK cells, *Lc*PPO, *Pd*CO, *Tr*PPO1, and *As*OPN3 showed the strongest expression and membrane targeting, and were only outperformed by *Dr*PPO1 (Fig. 1k). We selected the seven best-performing variants (across all assays; Extended Data Fig. 4) and next tested their ability to attenuate synaptic transmission.

### Benchmarking of bistable optoGPCRs in autaptic neurons

We expressed each of the selected optoGPCRs via rAAV2/1 transduction in autaptic neurons, a preparation in which single cultured neurons, grown on pre-seeded astrocyte micro-islands, form autaptic connections (Fig. 2a). Using whole-cell patch-clamp electrophysiology, we applied a series of paired pulses (pairs of depolarizations from -70 to 0 mV, every 5 s). We measured the induced excitatory postsynaptic current (EPSC) over three sequential periods: in the dark, post-light-application (one 500 ms pulse, 390 nm), and after administration of a 4.5 s-long 560 nm light pulse to inactivate the optoGPCR (Fig. 2b and Extended Data Fig. 5). We applied the same protocol to non-expressing controls, to correct for spontaneous EPSC rundown over time (Fig. 2c,d and Extended Data Fig. 5). Light activation of the PPOs from pufferfish (*Tr*PPO1/2) and zebrafish (*Dr*PPO1) had no effect on synaptic transmission, while *Ol*TMT1A, *Pd*CO, *Lc*PPO and *As*OPN3 significantly attenuated the AP-evoked EPSCs (Fig. 2e and Extended Data Fig. 5). *Ol*TMT1A and *Lc*PPO could only attenuate transmission by 66±5% and 61±5%, respectively. Activation of *Pd*CO and *As*OPN3 yielded the strongest EPSC reduction, by 89±3% and 84±5%, respectively (Fig. 2e). As reported previously^13^, *AsOPN3-*mediated inhibition was long-lasting and could not be recovered by light application at different wavelengths. Green light-induced EPSC recovery was only partially possible for *Ol*TMT1A, due to overlapping spectra of the opsin’s dark-adapted and active states^29^. However, for *Pd*CO and *Lc*PPO, the green light pulse reliably induced recovery of synaptic transmission (Fig. 2e).

**Figure 2:**
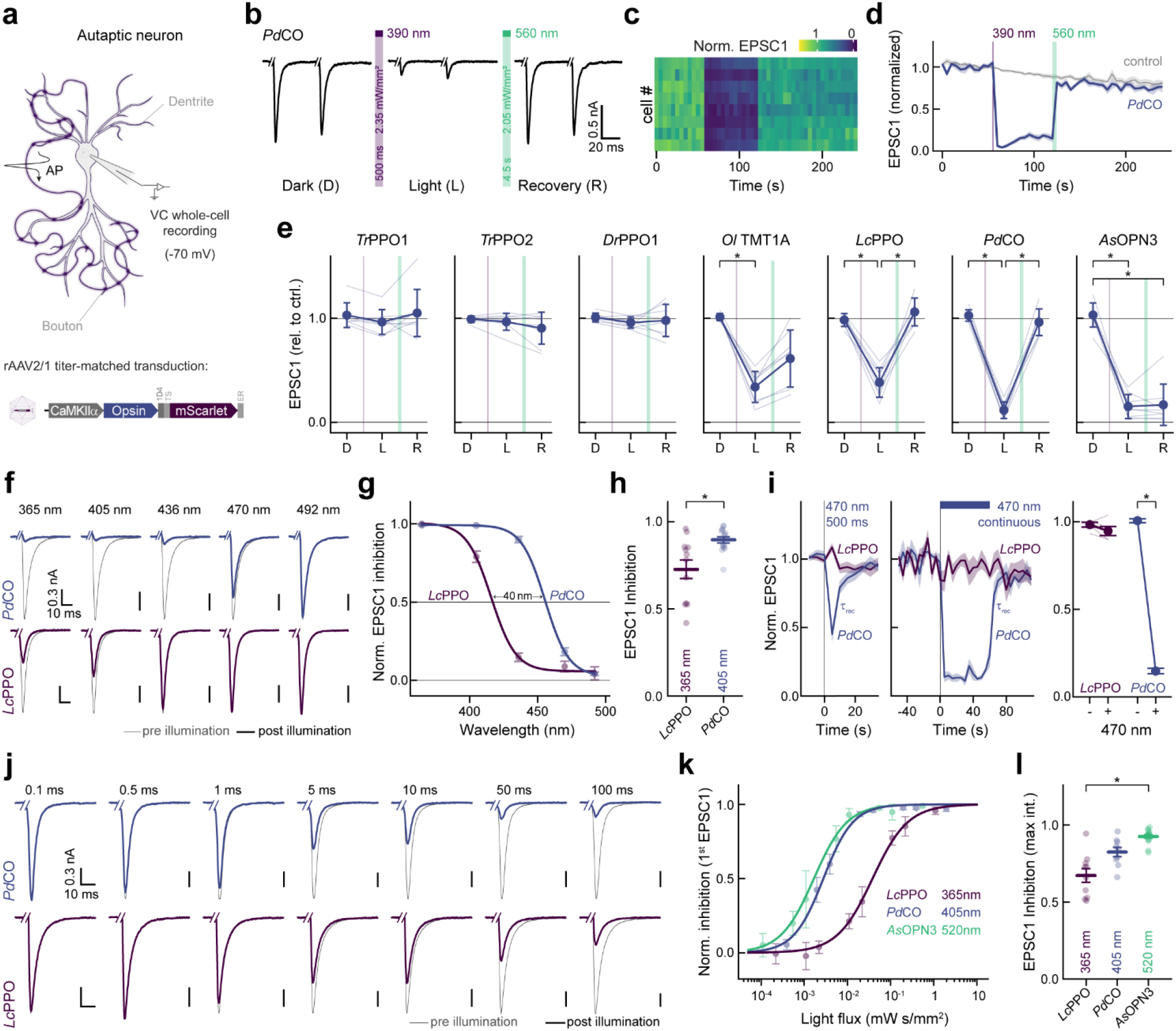
Comparison of optoGPCR performance and biophysical characterization in autaptic neurons. **a**, Schematic representation of an autaptic neuron recorded in the whole-cell configuration. Autaptic neurons were depolarized to fire unclamped action potentials, which triggered excitatory postsynaptic potentials (EPSCs). Neurons were transduced with rAAVs (bottom) encoding different optoGPCRs. **b**, Representative EPSCs traces evoked by a pair of 1 ms depolarizing current injections (50 ms inter-stimulus interval, every 5 s) in a *Pd*CO-expressing autaptic neuron before (left, Dark) and after (middle, Light) UV-light illumination, followed by green light-induced recovery (right, Recovery). Traces show 5 averaged sweeps. Current injection transients were removed for visualization. **c**, Contour plot representing EPSC amplitudes in 8 neurons expressing *Pd*CO, activated with 390 nm light for 500 ms and recovered by 560 nm light for 4.5 s. EPSCs were normalized to the average amplitudes of 5 EPSCs prior to 390 nm illumination. **d**, Averaged EPSC data (across replicates, n = 8) as shown in c, together with control EPSC recordings from non-expressing autaptic cultures measured with the same protocol. **e**, Quantification of EPSC inhibition for all optoGPCR candidates as shown in b-d. For dark (D; 2 EPSCs prior to UV light), light (L; 2 EPSCs post UV light and 2 EPSCs prior to green light) and recovery (R; 2 EPSCs 20s after green light) were averaged and normalized for EPSC rundown by the same quantification of non-expressing control cells from matching autaptic cultures as shown in c. Statistics: * indicates significance p<0.05; Friedman test followed by the Dunn-Sidak multiple comparison; *Ol*TMT1A: p(D,L)= 3.46E-03, p(D,R) 1.78E-02; *Lc*PPO: p(D,L)= 3.68E-02, p(L,R)= 1.40E-03; *Pd*CO: p(D,L)= 1.40E-03, p(L,R)= 3.68E-02; *As*OPN3: p(D,L)= 3.46E-03, p(D,R)= 1.78E-02. Data is shown as mean ± SD, n = 8. **f**, Example EPSCs (average of 7) pre illumination (gray) and post illumination (blue and purple), activated with light of different wavelengths (equal photon flux density) as indicated for *Pd*CO (top) and *Lc*PPO (bottom) from the same autaptic neuron respectively. **g**, Quantification of normalized EPSC inhibition for different wavelengths. For each cell, EPSC inhibition at each wavelength was normalized to the maximum inhibition. Lines show dose-response fit (n = 6 - 15). **h**, Quantification of the absolute EPSC inhibition at indicated wavelengths for experiments shown in f,g (n = 12 - 15). **i**, EPSC inhibition by blue light (470 nm) for *Pd*CO and *Lc*PPO. Illumination with blue light for 0.5 s (left, n = 10 - 14) or 60 s (middle, n = 5) evoked EPSCs inhibition by *Pd*CO but not *Lc*PPO. EPSCs recovered spontaneously for *Pd*CO with a time constant τ_rec_ (monoexponential fit, see main text). (right) quantification of the EPSC reduction by sustained application of blue light, n= 5. Statistics: * indicates significance p<0.05; two-sided Wilcoxon Rank Sum Test; p= 6.27E-03. **j**, Representative EPSCs traces (average of 7) pre (gray) and post illumination with different light pulse durations (blue and purple) recorded in the same autaptic neuron, for *Pd*CO (top) and *Lc*PPO (bottom). **k**, Quantification of release inhibition post illumination versus light flux for *Pd*CO, *Lc*PPO and *As*OPN3, normalized to the inhibition for maximum light flux used. Solid lines show sigmoidal fits (n = 3 - 17) **l**, Quantification of the absolute EPSC inhibition over 30 s post illumination at indicated wavelengths and maximum light flux for experiments as shown in j,k (n = 9 - 17). Statistics: * indicates significance p<0.05; Kruskal-Wallis test followed by Dunn-Sidak multiple comparison; p(*Lc*PPO, *As*OPN3)= 1.82E-04. If not stated otherwise all data is shown as mean ± SEM.

In line with presynaptic inhibition, the frequency but not the amplitude of miniature EPSCs (mEPSCs) was reduced for the four optoGPCRs that showed light-induced EPSC attenuation (Extended Data Fig. 6). We observed an increased paired-pulse ratio following opsin activation, which was most strongly pronounced for *Pd*CO and *As*OPN3, providing further support for their presynaptic site of action (Extended Data Fig. 6).

### Biophysical activation properties of PdCO

Given its promising performance in autaptic neurons and its photochromic properties, we characterized *Pd*CO’s biophysical properties in further detail in the context of synaptic inhibition and compared it with *Lc*PPO and *As*OPN3. First, we varied the wavelength of the activating 500 ms light pulse to generate action spectra for opsin activation (Fig. 2f). The wavelength needed for half maximum EPSC inhibition of *Pd*CO was red-shifted by 40 nm compared to *Lc*PPO (Fig. 2g and Extended Data Fig. 7). In addition, synaptic transmission at this wavelength range was more effectively reduced by *Pd*CO compared to *Lc*PPO (Fig. 2h). *Pd*CO activation with blue light (470 nm) showed transient inhibition that recovered with a time constant τ_rec_ of 3.4±0.6 s (Fig. 2i). We next tested if continuous blue-light illumination of *Pd*CO can evoke sustained inhibition that would similarly recover in the dark, without the need for a green pulse for inactivation. Indeed, continuous 470 nm illumination (2.83 mW/mm²) for 60 s reduced EPSCs by 85±1% in *Pd*CO-expressing neurons. Evoked EPSCs recovered spontaneously after the cessation of light, with a time constant of 2.7±0.3 s. In contrast, we were not able to achieve inhibition with 470 nm light for *Lc*PPO (Fig. 2i).

Next, we varied the light pulse duration at the maximal effective wavelengths to compare the light sensitivity of *Pd*CO, *Lc*PPO and *As*OPN3 (Fig. 2j). When quantifying the first EPSC after light activation, *Pd*CO (EC_50_= 3.1±0.4 µW s/mm²) showed similar sensitivity to *As*OPN3 (EC_50_= 1.9±0.3 µW s/mm², p=0.3217), whereas *Lc*PPO showed lower sensitivity with an EC_50_ of 30±2 µW s/mm² (Fig. 2k and Extended Data Fig. 7). At the maximum pulse duration, *As*OPN3 showed the strongest inhibition of (93±1%) followed by *Pd*CO (82±3%) and *Lc*PPO (67±4%; Fig. 2l).

We next tested whether *Pd*CO can be activated with two-photon excitation in HEK293T cells co-expressing GIRK channels. To obtain the two-photon action spectrum for *Pd*CO, we measured GIRK channel activation in cells expressing *Pd*CO using whole-cell patch-clamp electrophysiology (Extended Data Fig. 7d). First, we applied raster scans at different wavelengths ranging from 700 nm to 1100 nm (3 mW, 20 s raster scanning) while applying a voltage ramp from -120 mV to +40 mV. Maximum GIRK channel activation was achieved with 800 nm at -120 mV (Extended Data Fig. 7e), in good agreement with one-photon activation. Next, we titrated the *Pd*CO-coupled GIRK activation at 800 nm by varying light intensity. The half-maximum activation was determined to be 0.49±0.2 mW (Extended Data Fig. 7f).

### G-protein specificity

During the electrophysiological recordings in autaptic neurons, we observed stronger and faster GIRK-mediated hyperpolarization in neurons expressing *Pd*CO as compared to *As*OPN3 or *Lc*PPO (Fig 3a-d). To test whether the light-induced currents were caused by GIRK-channel coupling, we inhibited GIRK-channels with SCH23990 (Fig. 3a). Light-evoked GIRK-mediated currents were reduced by 77±6% after SCH23990 application in *Pd*CO-expressing neurons, while inhibition of EPSCs by *Pd*CO was not affected (Fig. 3e,f), indicating that synaptic inhibition via *Pd*CO is independent of GIRK channel activity. Because *Pd*CO showed very weak coupling to Gα_o_ and Gα_z_ in the GsX assay (Extended Data Fig. 2), we speculated that *Pd*CO might have a different G-protein signaling bias that leads to the observed differences in GIRK activation. We therefore employed the TRUPATH assay^30^ to characterize the G-protein signaling of *Lc*PPO, *As*OPN3 and *Pd*CO in more detail (Fig. 3g). Both *As*OPN3 and *Lc*PPO showed long-lasting coupling to all members of the inhibitory G-protein family (Gα_i-z_). In contrast, *Pd*CO only coupled to Gα_oA/B_ and Gα_z_, and not to Gα_i1-3_ (Fig. 3h and Extended Data Fig. 8a). To exclude that inhibition of synaptic transmission is mediated by activation of Gα_z_, we inhibited Gα_i_ and Gα_o_ coupling with pertussis toxin (PTX). For all three opsins, PTX treatment abolished light-induced inhibition of EPSCs, indicating that Gα_z_ does not contribute to presynaptic inhibition by these opsins (Fig. 3i and Extended Data Fig. 8b). As Gα_i_ proteins are the main inhibitors of adenylate cyclases, we tested whether *Pd*CO is capable of modulating cAMP production using a cAMP-dependent luciferase assay (GloSensor; Fig. 3j). As anticipated, *Pd*CO activation did not have any detectable effect on cAMP production, whereas *Lc*PPO and *As*OPN3 activation led to a bioluminescence signal decrease of 63±1% and 62±1%, respectively (Fig. 3k,l).

**Figure 3:**
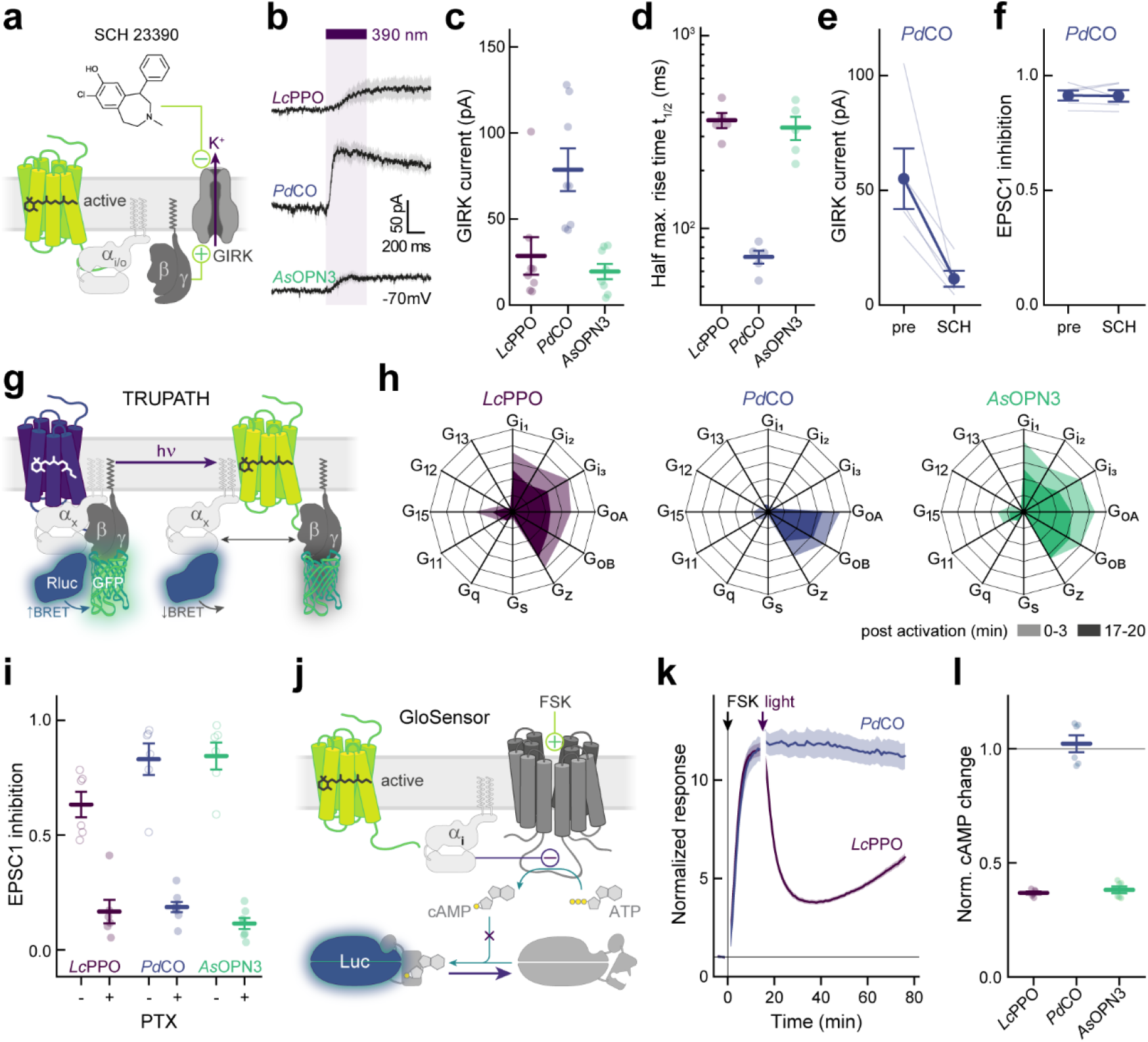
G-protein signaling in potent inhibitory optoGPCRs. **a**, Scheme of GIRK activation and inhibition. Gβγ-complex signaling by photoactivated optoGPCRs activates GIRK channels, which are inhibited by SCH23390. **b**, Average current traces from cells expressing the indicating optoGPCRs, showing GIRK channel activation induced by light application in autaptic neurons. **c**, Quantification of maximum GIRK channel currents (n = 8). **d**, Quantification of the rise time to the half maximum current determined from the neurons shown in b. **e**, GIRK channel currents activated by light and inhibited by local application of SCH23390 in autaptic neurons (n = 5). **f**, EPSC1 inhibition of the same neurons shown in e. Application of SCH23390 does not alter EPSC inhibition. **g**, TRUPATH assay. Activation of the optoGPCR reduces the bioluminescence energy transfer (BRET) between the luciferase-fused Gα and the GFP-fused Gβγ subunits. **h**, Relative -netBRET means (n = 4) for major Gα subtypes integrated for 0-3 min post light activation (1s 390 nm; light shaded area) and integrated for 17-20 min post light activation (dark shaded area). **i**, EPSC1 inhibition as measured in Fig. 2a-e in autaptic neurons (open circles) and pertussis toxin (PTX) treated neurons from matched neuronal cultures. Cells were incubated with PTX for 12-16 h (n = 6 - 8). **j**, The GloSensor assay for inhibition of adenylyl cyclase (AC) activity by optoGPCRs. Forskolin (FSK) stimulates cAMP production by ACs, which can be inhibited by Gαi signaling induced by optoGPCRs. Changes in cAMP levels can be detected by cAMP-dependent Luciferase (Luc). **k**, Time course of cAMP response measured with the GloSensor assay for *Pd*CO and *Lc*PPO, after addition of FSK (t = 0 min; black arrow) and after activation of *Lc*PPO and *Pd*CO (purple arrow). Data is normalized to bioluminescence reads pre-FSK application (n = 6). **l**, Normalized cAMP changes after light application, calculated by division of minimum response post illumination by maximum pre-illumination response of data as shown in k (n = 6). All data is shown as mean ± SEM.

### Presynaptic inhibition in organotypic hippocampal slices

We next aimed to assay the inhibition efficacy of *Pd*CO against *Lc*PPO, the only other photoswitchable optoGPCR using organotypic hippocampal slice cultures. First, we confirmed that the biophysical properties of these two opsins were similar to those characterized in the autaptic culture preparation. Individual CA3 pyramidal neurons were transfected by single-cell electroporation to express either *Pd*CO (Extended Data Fig. 9a) or *Lc*PPO. We recorded GIRK-mediated currents evoked by light pulses at varying wavelengths and durations (Fig. 4a). The maximum GIRK current response for *Pd*CO-expressing neurons was between 405 nm and 435 nm (Fig. 4b). Peak GIRK currents evoked by *Pd*CO were higher than the ones induced by *Lc*PPO at all tested wavelengths, even at a 10-fold lower light intensity for *Pd*CO. Next, we varied the illumination time at the peak activation wavelengths of both optoGPCRs (365 nm for *Lc*PPO and 405 nm for *Pd*CO). *Pd*CO-evoked GIRK currents showed maximum responses to light pulses with durations between 50 and 100 ms, and a higher amplitude than those evoked by *Lc*PPO at the same pulse duration (Fig. 4c). We next activated the two optoGPCRs selectively at axonal terminals (Extended Data Fig. 9b) to compare their ability to suppress synaptic transmission. Presynaptic CA3 neurons were virally co-transduced with *Pd*CO or *Lc*PPO together with a soma-localized BIPOLES (somBIPOLES)^31^, to elicit red light (625 nm)-evoked APs in CA3 while avoiding potential cross-activation by *Pd*CO illumination at CA1 (Fig. 4d). Red light pulses applied to the CA3 region reliably evoked EPSCs in CA1 cells, while application of 100 ms light pulse at 365 nm (10 mW/mm²) to *Lc*PPO-expressing terminals reduced EPCS by 27±4% (Fig. 4e,f and Extended Data Fig. 9). Activation of *Pd*CO with 10-fold lower light power at 405 nm led to a 78±5 % reduction in synaptic transmission, while no EPSC reduction was observed when somBIPOLES was expressed alone (Fig. 4e,f and Extended Data Fig. 9). For both optoGPCRs, attenuation of synaptic transmission was reliably recovered with 525 nm light (Fig. 4e,f).

**Figure 4:**
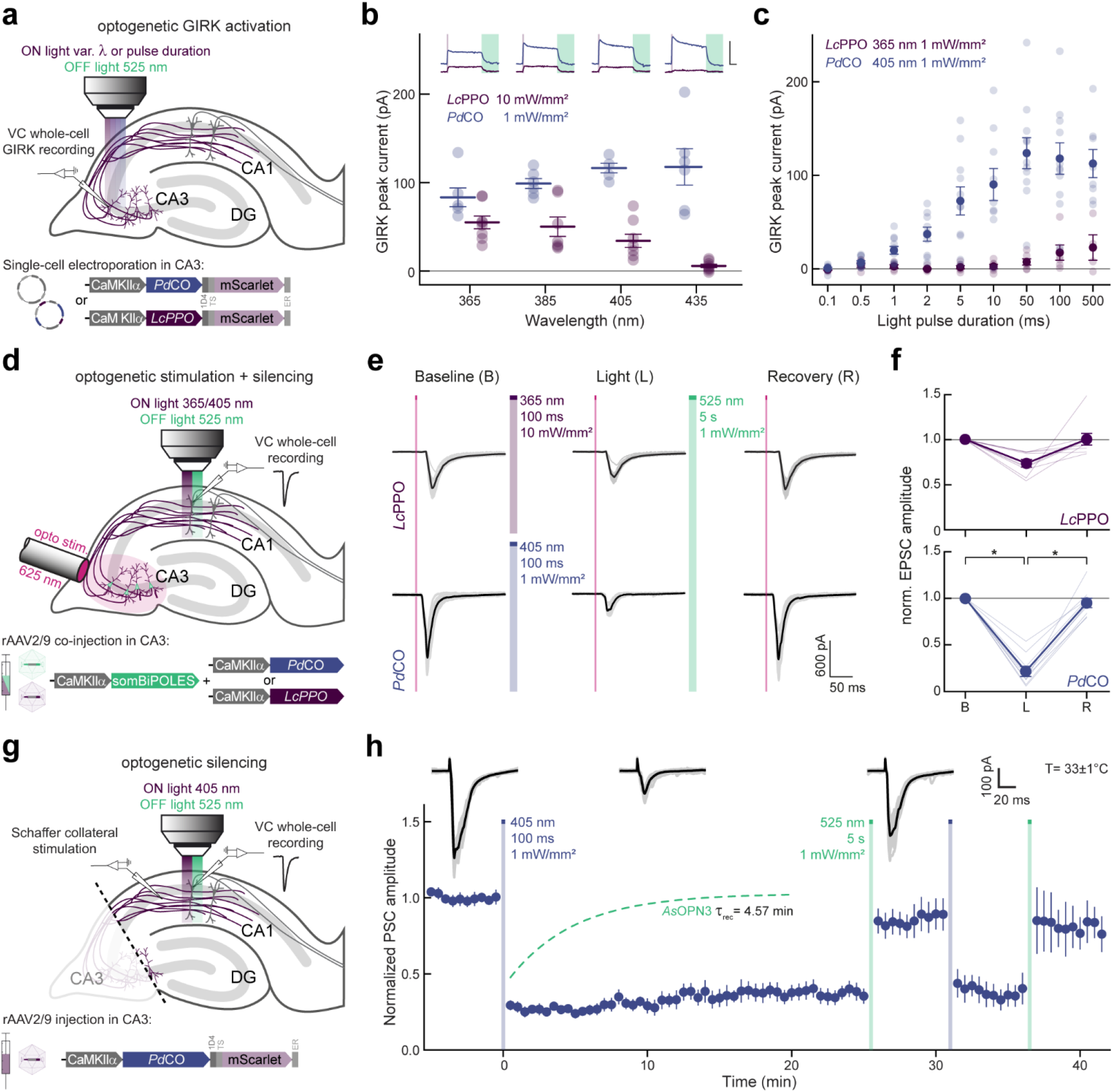
Characterization and performance of optoGPCRs in organotypic hippocampal slice cultures. **a**, Illustration of experimental setup in organotypic hippocampal slice cultures (single cell plasmid electroporation, circles). Activation spectrum and light sensitivity were measured by recording GIRK-mediated currents from CA3 neurons expressing either *Pd*CO or *Lc*PPO in response to optogenetic stimulation through the microscope’s objective at varying light parameters. **b**, Quantification of GIRK current amplitudes recorded in cells expressing *Pd*CO and *Lc*PPO at the indicated light wavelengths and intensities. Activation light pulse: 500 ms. Inactivation light pulse: 5 s at 525 nm. Inset scale bar: 100 pA, 2 s. n = 6 - 8 **c**, Quantification of GIRK current amplitudes recorded in cells expressing *Pd*CO and *Lc*PPO at their optimal activation wavelength at varying light durations (n = 4 - 14). **d**, Illustration of experimental design for bidirectional optogenetic control of synaptic transmission. **e**, Representative current traces of patched CA1 neurons in response to optogenetically-induced presynaptic APs (625 nm, 5 ms) under baseline conditions (left), after activation (middle) and inactivation (right) of the optoGPCRs. Top: *Lc*PPO group. Bottom: *Pd*CO group. Gray: single trials; black: averaged traces. **f**, Normalized EPSC amplitudes from *Lc*PPO group (top) and *Pd*CO group (bottom). n = 9. Statistics: * indicates significance p<0.05; Repeated measures one-way ANOVA with the Geisser-Greenhouse correction followed by Tukey’s comparison; PdCO: p(B,L)< 1.00E-04, p(L,R)= 1.00E-04. **g**, Illustration of experimental setup for electrical stimulation of Schaffer collaterals and optogenetic inhibition of synaptic transmission. **h**, Time course of normalized PSC amplitudes from all the recorded postsynaptic CA1 neurons, before, during and after activation/inactivation of *Pd*CO. Representative voltage-clamp traces are shown on top. Gray: single trials; black: average trials. Light was applied locally in CA1 for activation and inactivation of *Pd*CO (ON light: 500ms, 405 nm, 1 mW/mm^2^; OFF light: 5 s, 525 nm, 1 mW/mm^2^). n= 3 - 7. All data is shown as mean ± SEM.

We next measured the stability of inhibition of synaptic release by *Pd*CO, by stimulating *Pd*CO-expressing Shaffer collaterals with a bipolar electrode at 0.1 Hz, while recording EPSCs in CA1 neurons (Fig. 4g). To exclude any somatic effects of the opsin and to avoid antidromic and recurrent activation of the CA3 network, we dissected out area CA3 prior to the recordings.

Local application of a brief 500 ms light pulse in CA1 reduced evoked PSCs by 71 ± 0.3% and showed no spontaneous recovery over the time course of 25 minutes (Fig. 4h). This is in contrast to *As*OPN3-mediated inhibition, which spontaneously recovers with a time constant of approximately 5 minutes under identical experimental conditions^13^. In addition, we were able to recover transmission with 525 nm light and subsequently block synaptic transmission again with a second 405 nm pulse (Fig. 4h). Normalized EPSC amplitudes were not affected in non-expressing control cultures or non-illuminated *Pd*CO cultures. (Extended Data Fig. 9).

### Single-photon spectral multiplexing with PdCO

It is often informative to combine an optical readout of neuronal activity with optogenetic manipulations. For example, fiber photometry or, more recently, miniature microscopes can be combined with light stimulation at a different wavelength in the single-photon domain^32,33^. This requires spectral multiplexing of different optogenetic sensors and actuators, and benefits from minimizing spectral crosstalk^1,34^. To establish whether *Pd*CO can be combined with red-shifted sensors or actuators, we next analyzed the wavelength-dependence of inactivation by varying the wavelength and irradiance of the inactivating pulse for both *Pd*CO and *Lc*PPO expressed in autaptic neurons. In these experiments, the optoGPCRs were activated at their peak excitation wavelength, and inactivation light was applied 30 s later (Fig. 5a). *Lc*PPO showed a broad wavelength sensitivity that enabled near complete off-switching between 436 and 560 nm, while *Pd*CO’s inactivation sensitivity was maximal between 470 and 520 nm (Fig. 5b and Extended Data Fig. 2b). We noted that the confined spectral window for inactivating *Pd*CO might present an opportunity for spectral multiplexing with other optogenetic probes that are activated by longer wavelengths. We therefore titrated the light sensitivity for both optoGPCRs at 560 nm and determined that EPSC recovery at this wavelength is 6-fold more efficient for *Lc*PPO (EC_50_= 61±2 µW/mm²) compared to *Pd*CO (EC_50_ = 372±163 µW/mm²; Fig. 5c), suggesting that *Pd*CO is better suited for multiplexing applications with red-shifted sensors or actuators.

**Figure 5:**
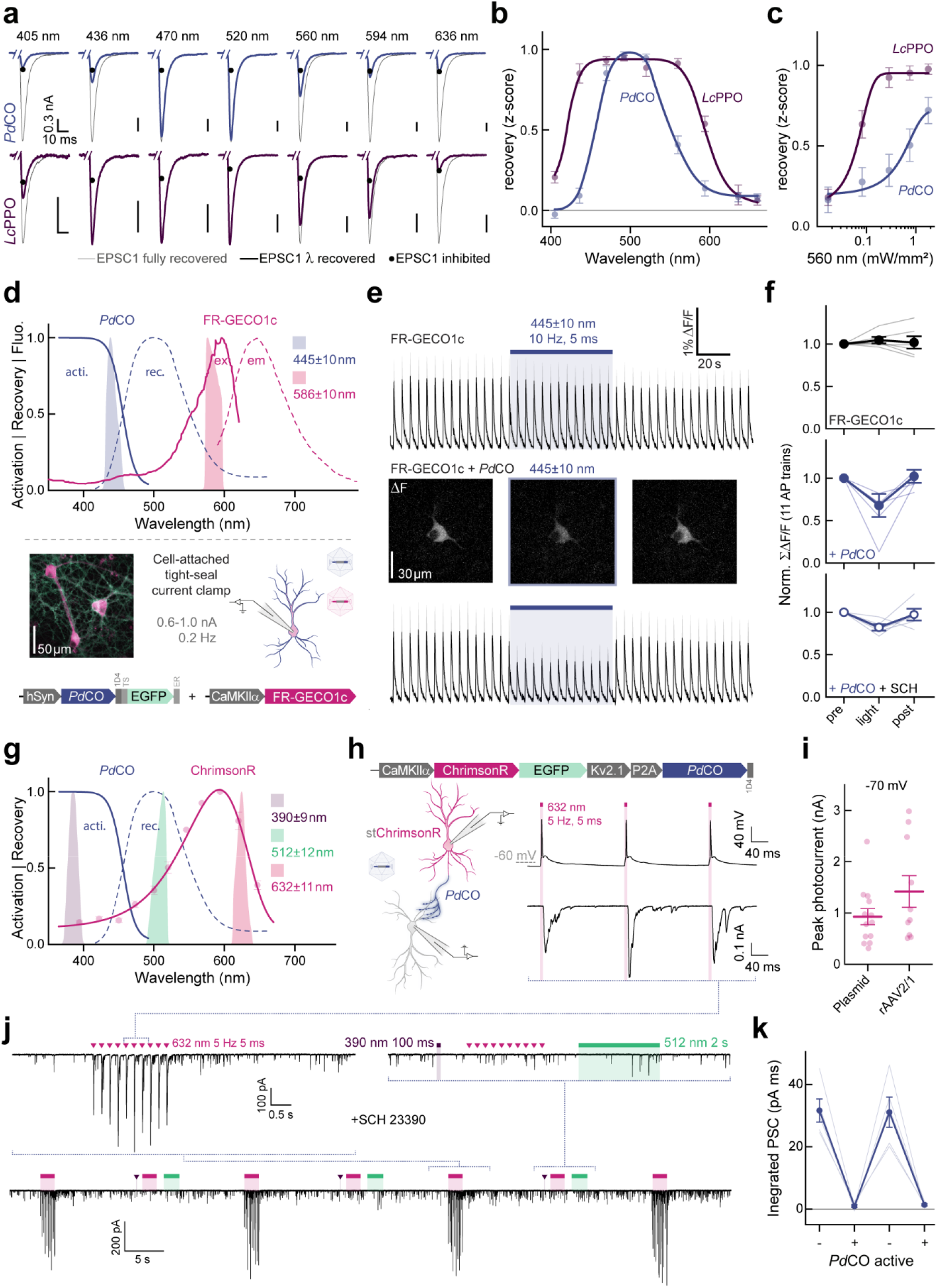
Single-photon spectral multiplexing of *Pd*CO. **a**, Example traces of experiments used to determine the spectral features and light sensitivity of optoGPCR inactivation in autaptic neurons. Samples were first illuminated with 390 nm (*Lc*PPO) or 405 nm (*Pd*CO) light for 500 ms, respectively, to inhibit EPSCs (black circles), followed by recovery with light delivery at the indicated wavelengths (equal photon flux density) for 4.5 s (colored traces) and finally completely recovered with 520 nm for at least 10s (gray traces). An average of 7 EPSC traces are shown, scaled to the fully recovered EPSC. **b**, Wavelength sensitivity of light-induced recovery. To correct for potential EPSC rundown, recovery z-scores were calculated using the mean of 4 EPSC post inhibition, prior recovery light at different wavelengths and the mean of 4 EPSC post full recovery with green light. n = 4 - 7. **c**, Light titration of light induced recovery. Experiments were conducted as in b but at a fixed wavelength of 560 nm, while varying the light intensity between trials. n = 7 - 8. **d**, Top: spectra of *Pd*CO activation (solid blue line) and inactivation (dashed blue line), excitation (solid magenta line) and emission (dashed magenta line) spectra of FR-GECO1c^40^ and stimulation light properties used for activation/ excitation (blue/ magentas shaded areas). Bottom (upper left): representative epifluorescence pseudocolor image of neuronal culture transduced with AAVs encoding *Pd*CO-EGFP and FR-GECO1c. Bottom (upper right): schematic approach of the cell-attached tight-seal patch-clamp configuration. Bottom: constructs used for spectral multiplexing calcium imaging experiments. **e**, Average (across replicates) FR-GECO1c calcium traces during repetitive current injections (top) and with additionally expressing *Pd*CO (lower trace). Example pictures from left to right: averaged signal before, during, and after blue light illumination. **f**, Quantification FR-GECO1c only (Top), co-expressed with *Pd*CO (middle) and (bottom) additionally blocking GIRK currents with SCH23390 (SCH). n = 5 - 6. **g**, Spectra of *Pd*CO activation (solid blue line) and inactivation (dashed blue line), together with the action spectra recorded from stChrimsonR-EGFP-P2A-*Pd*CO (see e) expressing neurons measured with TTX, CNQX, AP-5 at -70 mV holding potential. Action spectra were recorded twice per cell in both directions (UV to red and vice versa) with equal photon flux density (2 ms light pulse) and then averaged. Stimulation light properties (purple/green/magenta shaded areas) are shown. n = 7. **h**, Construct design (top), schematic representation of expression (bottom left) and experiment (bottom right) are shown. Red-light activation (632 nm, 5 ms, 5 Hz) evoked APs in stChrimsonR-EGFP-P2A-*Pd*CO expressing cells (upper trace), while in non-expressing cells a pronounced PSC could be recorded (lower trace). **i**, Quantification of maximum photocurrent amplitudes in stChrimsonR-EGFP-P2A-*Pd*CO expressing cells mediated by either calcium-phosphate transfection or viral transduction. n = 10 - 13. **j**, Upper left trace: Representative recording of a postsynaptic non-expressing cell, where red light application evoked reliable PSCs. Upper right trace: The same neuron as before but UV-light activation of *Pd*CO prior to ChrimsonR activation completely abolishes generation of PSCs. Bottom trace: Full time scale of the experiment showing repetitive suppression and recovery of ChrimsonR induced PSCs by toggling the activity of PdCO by UV and green light. **k**, Quantification of experiments shown in g for 5 biological replicates. All data is shown as mean ± SEM.

We next explored spectral multiplexing using the red-shifted calcium indicator FR-GECO1c^35^ (Fig. 5d, top). *Pd*CO, fused to EGFP for verification of expression, was co-expressed with FR-GECO1c in cultured neurons. In a tight-seal cell-attached patch-clamp configuration, we evoked APs at 0.2 Hz (Fig. 5d, bottom and Extended Data Fig. 10), resulting in reliable calcium transients in the FR-GECO1c signal (Fig. 5e, top). Blue light (445±10 nm) used to transiently activate *Pd*CO, caused a 32±14% reduction in the amplitude of evoked calcium events (Fig. 5e,f). Notably, as reported for other GECO variants^36^, blue light application alone increased FR-GECO1c fluorescence, for which we corrected in our analysis (Extended Data Fig. 10). As GIRK activation can lead to reduced excitability or even suppression of AP firing, we blocked GIRK channels using SCH23390. Blue light application still decreased calcium transients by 18±4%, while in control cells only expressing FR-GECO1c, no reduction of calcium transients was detected (Fig 5e,f). Consistent with previous work^13,14^, this indicates that *Pd*CO activation leads to the attenuation of somatodendritic VGCC activity.

Next, we combined *Pd*CO with a soma-targeted variant of the red light-sensitive channelrhodopsin ChrimsonR^37^ in a single bicistronic construct to allow the triggering of APs with red light, while simultaneously inhibiting synaptic transmission with the blue light-sensitive *Pd*CO (Fig. 5 g-k). In cultured hippocampal neurons expressing this bicistronic construct, red light pulses (5 ms, 632 nm) generated photocurrents above 900 pA that reliably induced APs (Fig. 5 h,i). In non-expressing neurons, the same red light pulses caused reliable postsynaptic currents (PSCs) (Fig. 5h, bottom trace). When activating *Pd*CO by a brief 390 nm light pulse (100 ms), repeated red light pulsing did not evoke PSCs anymore, indicating effective *Pd*CO-mediated inhibition of synaptic transmission (Fig. 5j, upper left trace). Following green light application (512 nm) to recover transmission, red light-evoked PSCs were readily detectable again (Fig. 5j, lower trace); inhibition of evoked synaptic transmission could be achieved in a repetitive manner (Fig. 5j,k).

### PdCO applications in cardiomyocytes and in vivo

To establish the efficacy of *Pd*CO in modulating mouse behavior, we used it to unilaterally inhibit dopaminergic projections from the substantia nigra to the dorsomedial striatum, a neural pathway known to play an important role in animal locomotion^38^. We activated *Pd*CO unilaterally in these axons during free locomotion and measured the resulting side bias, a behavioral test we previously established^13^. We expressed *Pd*CO (or EYFP as control) bilaterally in substantia nigra pars compacta (SNc) dopaminergic neurons and implanted bilateral optical fibers above the dorsomedial striatum (DMS; Fig. 6a). Unilateral light activation caused an ipsiversive rotational bias in *Pd*CO-expressing mice (Fig. 6b) that accumulated over time and ceased after illumination with green light (Fig. 6c.). This effect was consistent across *Pd*CO expressing mice and absent in the EYFP-expressing control group (Fig. 6c,d).

**Figure 6:**
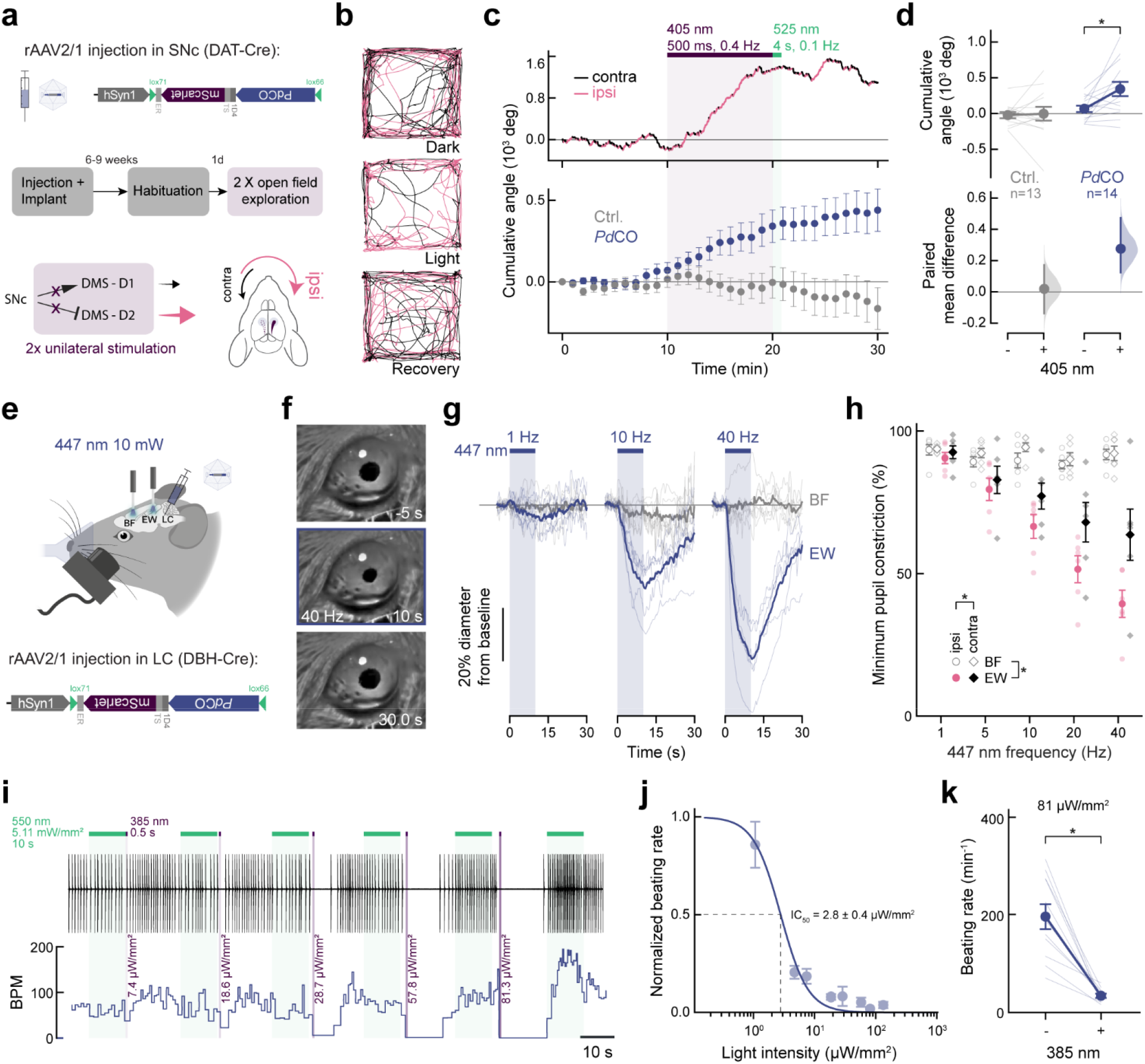
*Pd*CO applications in cardiomyocytes and in vivo. **a**, Schematics of the experimental setup and hypothesis. Top: schematic of the *Pd*CO construct. Middle: experimental timeline. Bottom: Bilateral expression of *Pd*CO in SNc dopaminergic neurons and unilateral light-mediated suppression of their striatal projections would induce an ipsiversive side bias during free locomotion. **b**, Locomotion trajectories of representative *Pd*CO mice, over successive 10-min periods: (top to bottom) before, during and after light delivery. Magenta and black color code ipsilateral or contralateral angle trajectory segments, respectively. **c**, Top: Representative cumulative angle traces of the individual *Pd*CO-expressing mice shown in e, over 30 min of free locomotion in an open field arena. Magenta and black colors depict ipsilateral or contralateral segments, respectively. Bottom: Average cumulative angle across *Pd*CO-expressing (blue) and EYFP-expressing control mice (gray). Each mouse underwent two unilateral stimulations of each hemisphere, respectively, that was then averaged per mouse. n= 13-14 **d**, Quantification of the accumulated angle prior illumination (min. 9) compared to post UV-illumination (min. 20) for *Pd*CO-expressing (blue, n=14) and EYFP-expressing control mice (gray, n= 13). The paired mean difference for both comparisons is shown as Cumming estimation plot. Each paired mean difference is plotted as a bootstrap sampling distribution. Mean differences are depicted as dots; 95% confidence intervals are indicated by the ends of the vertical error bars. Statistics: * indicates significance p<0.05; two-sided permutation t-test; 5.80E-03. **e**, Schematic of pupil experiment (see methods). **f**, Representative frames from pupil video recording at t=-5 (top),10 (middle) and 30 s (bottom) relative to 40 Hz laser stimulation onset. **g**, Mean pupillometry traces (bold lines, n=6) for 1 Hz (left), 10 Hz (middle), and 40 Hz (right) laser stimulation. Each time course denotes the median time-course of ipsilateral pupil diameter across trials in each subject in the basal forebrain (BF, gray) and the Edinger-Westphal nucleus (EW, blue). The vertical blue shaded area represents laser stimulation interval. Thin traces, individual mice; Thick traces, mean (n=6). **h**, Average pupil constriction (n=6 mice, matching the minimal value in time-courses shown in panel j) as a function of stimulation frequency (x-axis). Magenta dots and black diamonds represent EW stimulation effect on ipsilateral pupil or contralateral pupil, respectively. Bold dots/diamonds, average across mice (n=6). Error bars, SEM across mice. Light dots, average of trials in each mouse. Note that EW stimulation is associated with dose-dependent pupil constriction, and this is stronger for ipsilateral pupils. Statistics: * indicates significance p<0.05; multiple way ANOVA; p(placement)= 1.22E-20, p(frequency)= 9.62E-12, p(eye laterality)= 1.98E-04, p(placement·frequency)= 5.42E-10, p(placement·eye laterality)= 8.71E-03. **i**, Top: representative beating trace (one dash per beat) of an atrial cardiomyocyte expressing *Pd*CO, activated with different UV-light intensities and inactivated with green light. Bottom: Analysis of beats per minute (BPM) over time. **j**, Quantification of normalized beating frequency vs applied UV-light intensities from n = x different cells. Solid line depicts dose-response fit used for EC_50_ estimation. **k**, Quantification of beating reduction at 81 µW/mm^2^ UV-light compared to non-illuminated conditions. Statistics: * indicates significance p<0.05; two-tailed paired t-test; p= 5.35E-05. If not stated otherwise all data is shown as mean ± SEM.

To further test how *Pd*CO inhibits specific synapses *in vivo*, we focused on locus coeruleus norepinephrine (LC-NE) modulation of pupil size^39–41^, which is largely mediated by disinhibition of the parasympathetic Edinger-Westphal (EW) nucleus^42^. LC neurons form vast, widespread projections that terminate in multiple brain regions^43^, making somatic inhibition highly non-specific. To selectively suppress LC axons terminating in the EW nucleus, we conditionally expressed *Pd*CO unilaterally in NE neurons of the LC, and implanted optical fibers above the ipsilateral EW, and above the basal forebrain (BF, as control region; Fig 6e). Blue light application (447 nm) to EW led to robust dose-dependent pupil constriction, whereas identical stimulation of BF did not (Fig. 6f-h). Notably, the ipsilateral pupil was significantly more affected by laser stimulation than the contralateral pupil. Given that pupil asymmetry does not occur under physiological conditions^44^, the observed lateralization provides strong evidence for successful pathway-specific inhibition (Fig. 6h).

Finally, we investigated the potential application of *Pd*CO in non-neuronal cardiac tissue. In the heart, multiple GPCR-mediated mechanisms modulate pacemaking activity. Sympathetic stimulation is mediated via β-adrenergic Gα_s_-signaling while parasympathetic stimulation occurs through M2-muscarinic receptor Gα_i_-signaling. Activation of Gα_s_ or Gα_i_ increases or decreases AC activity and cAMP levels, respectively. Elevated cAMP levels activate hyperpolarization-activated cyclic nucleotide-gated (HCN) ion channels leading to faster depolarization and beating frequency. While *Pd*CO does not inhibit AC (Fig. 3j-l), it strongly couples to GIRK channels (Fig. 1g-i) and suppresses calcium transients (Fig. 5d-f). Since atrial cardiomyocytes endogenously express high levels of GIRK channels and rely on calcium influx for their electrical activity, we expressed *Pd*CO in spontaneous active neonatal atrial cardiomyocytes and explored light-mediated suppression of pacing. We found that the spontaneous beating of *Pd*CO-expressing cardiomyocytes could be suppressed after a brief violet light pulse in a light intensity-dependent manner (Fig. 6i,j). Inhibition of beating occurred with a half-maximal light dose of 2.8±0.4 µW/mm² (Fig. 6j), consistent with similar measurements in neurons. At 81 µW/mm², beating could be efficiently reduced by 80±4% (Fig. 6k). These results suggest that *Pd*CO can be efficiently utilized in non-neuronal tissues with comparable efficiency and light sensitivity to neuronal preparations.

## Discussion

Efficient presynaptic inhibition offers unique opportunities to study projection-specific contributions to behavior. While most previous optogenetic tools are not suitable for this approach^8,9,16,45^, two recently described optoGPCRs (*As*OPN3 and *Lc*PPO) could sufficiently provide axonal inhibition. However, each of these powerful tools has critical limitations that impair their utility for optogenetic applications, including a wide action spectrum and long spontaneous decay kinetics (*As*OPN3) and a broad and light-sensitive deactivation cross-section that limits multiplexing applications (*Lc*PPO). We therefore systematically compared a set of bistable type II rhodopsins with the goal of identifying optoGPCR candidates suitable for efficient presynaptic inhibition with precise temporal control and spectral properties suitable for optical multiplexing.

Both *Lc*PPO and *Pd*CO are photoswitchable optoGPCRs that can be activated with UV to blue light and inactivated with green light. For experiments requiring rapid switchability, these opsins have an inherent advantage over *As*OPN3, which is activated more broadly with any wavelength between 390 and 640 nm, and only recovers through a thermal back-reaction in the dark. Thus, *Pd*CO and *Lc*PPO can provide superior temporal control over termination of optoGPCR signaling. *Pd*CO showed faster onset compared to *As*OPN3 and *Lc*PPO, allowing a more precise timing of G-protein signaling onset. However, the rate limiting steps in signaling kinetics will be determined in all cases by the availability and mobility of the Gα and Gβγ subunits. Importantly, *Pd*CO showed more robust suppression of presynaptic transmission when compared to *Lc*PPO in hippocampal neurons, and its action spectrum is red-shifted by 40nm, allowing maximal activation with wavelengths up to 420nm. Blue light (>440nm) can be used to transiently activate PdCO since it is absorbed by both the inactive and active states and therefore drives both the forward- and back-reactions simultaneously.

When tested as a purified protein, the absorption maximum of *Pd*CO (383nm)^26^ is only shifted by 13nm compared to that of *Lc*PPO (370nm)^23^. However, we were able to activate *Pd*CO with wavelengths up to 470nm. For *Lc*PPO, synaptic inhibition was only feasible up to 405nm in our recordings, and GIRK activation in organotypic slices dropped strongly above 405nm, in line with previous reports^24^. *Pd*CO’s red-shifted activation spectra can be explained by a low absorption cross-section of the active state that shifts the equilibrium wavelength between activation and light-induced recovery towards the red spectrum. The lower probability of light-induced inactivation is also reflected in the inactivation spectrum of *Pd*CO, which is narrower compared to that of *Lc*PPO. Intriguingly, *Lc*PPO has been reported to be activatable with blue light (around 470 nm) in various experimental settings^14^. However, we were not able to achieve true blue light activation of *Lc*PPO. Instead, we efficiently inhibited *Lc*PPO activity in our experiments using 470 nm as reported previously^24^. This discrepancy might result from bandwidth limited light in our experiments, which eliminated low-wavelength photons.

We observed that *Pd*CO activation led to transient synaptic inhibition when low light intensities at 405 nm or more red-shifted activation light is used (e.g. 445-470 nm). This indicates that if a low number of the activated G-proteins are recruited, the inhibitory effects can be transient, consistent with similar effects in chemogenetic actuators^46^. Although we demonstrated synaptic inhibition over 20 mins in organotypic slice preparations, care should be taken when using *Pd*CO for long-lasting synaptic silencing experiments following only a brief single light pulse activation. Especially for *in vivo* experiments, if light delivery and expression levels are below saturation, *Pd*CO-mediated inhibition could be short-lived; this can be overcome by repetitive light application or increased opsin expression. Furthermore, optoGPCR kinetics might vary between cell types, availability of heterotrimeric G-protein subunits and effectors/targets^47^, and input specific AP frequency and membrane depolarization^48–50^. Therefore, the inhibitory effect of presynaptic optoGPCRs should be tested by recording postsynaptic input reduction over time as discussed elsewhere^16^. Such experiments would be facilitated by the bicistronic constructs described above (Fig. 5g-k), which would allow co-expression of the red-shifted ChrimsonR with *Pd*CO in the same neurons.

In our experiments, as well as across the optoGPCR literature, testing the same optoGPCR with different established assays of GPCR signaling can lead to vastly different outcomes. OPN5 homologs, for example, which did not couple to either Gα_q_ or Gα_i/o_ in the GsX assay. still generated a GIRK response. These optoGPCRs have recently been described to mediate Gα_q_ coupling in various settings^51,52^. For *Pd*CO, we observed efficient GIRK coupling as shown previously^26^ but could only demonstrate very weak Gα_o_ coupling in the GsX assay. The TRUPATH assay, however, revealed that *Pd*CO selectively couples to Gα_o_ but not to Gα_i,_ which was confirmed by the demonstration that *Pd*CO activation does not inhibit cAMP production. Nevertheless, we found that *Pd*CO allows efficient silencing of presynaptic transmission, indicating that selective activation of the Gα_o_ pathway can strongly suppress presynaptic release in all preparations tested in our study. The lack of *Pd*CO impact on AC activity and the absence of effects on presynaptic cAMP can offer potential benefits as cAMP is involved in various intracellular processes such as proliferation, differentiation, survival, long-term synaptic potentiation, neurogenesis, and neuronal plasticity. However, potential modulation of cAMP by *Pd*CO activation should not be completely excluded as a variety of ACs have been reported to be affected by different Gα and Gβγ subunits including Gα_o_ ^55^. It has also been shown that *Pd*CO can transiently recruit Gα_i_ under long lasting continuous and/or high-intensity illumination^53,54^, potentially by depleting available Gα_o_ over time and therefore generating a signaling bias towards Gα_i_.

While our primary focus in this study has been to develop new optoGPCRs for presynaptic inhibition, *Pd*CO and many other optoGPCRs showed strong coupling to GIRK channels at the neuronal soma. Since *Pd*CO showed both faster and stronger GIRK coupling in autaptic neurons compared to *As*OPN3 and *Lc*PPO, it could be used as a tool to reduce neuronal excitability when activated at the soma. However, not all neurons express GIRK channels and somatic inhibition might be absent in some cell types (e.g. medium spiny neurons in the striatum). Thus, when somatic inhibition is desired, anion- or potassium-conducting channelrhodopsins^10,55,56^ might be more suitable, due to their strong inhibitory photocurrents and their millisecond-scale decay kinetics upon light offset. Nonetheless, by blocking GIRK channel activity we demonstrated that *Pd*CO-mediated synaptic attenuation of transmission is independent of GIRK activity and can therefore be applied in neurons lacking these channels.

The diversity of genetically-encoded actuators and sensors provides a wealth of opportunities for multiplexed experiments, combining two or more such tools in a single experimental setting. As the activation spectrum of *As*OPN3 covers the entire UV-visible range and due to its high light sensitivity, it requires careful handling and cannot be combined with other optical approaches apart from two-photon imaging^13^. In contrast, *Lc*PPO and *Pd*CO are both activated on the high energy visible spectrum (UV to blue light) and therefore do not bear the risk of cross-activation by other wavelengths used for imaging or optogenetic control. The narrow action spectrum of *Pd*CO’s light-induced back-reaction to the inactive state is an attractive property for multiplexing with genetically-encoded tools that have red-shifted excitation spectra. Whereas one photon multiplexing has been demonstrated to be similarly possible with *Lc*PPO^14^, we found that application of cyan to red light can cause a stronger inactivation of *Lc*PPO compared to *Pd*CO (Fig. 5a-c). For activation of larger brain areas, *As*OPN3 might serve as a more suitable inhibitory optoGPCR due to its red-shifted activation spectrum and high light sensitivity. However, independent *As*OPN3 activation at different brain loci might be less feasible due to potential cross-excitation by scattered photons. In this case, *Lc*PPO and *Pd*CO could serve as an alternative as UV-blue light is more effectively attenuated in neuronal tissues. Since *As*OPN3 can also be activated by UV to blue light, these wavelengths can be used to excite *As*OPN3 in settings where slow kinetics are desirable and activation by scattered light is a concern.

Taken together, we demonstrated that *Pd*CO is a rapid, reversible, and versatile optoGPCR that mediates inhibition of synaptic transmission efficiently in diverse cell types *in vitro* and *in vivo*. With activation time constants in the sub 100-ms and inactivation switching times <10 s, *Pd*CO serves as a fast inhibitory optoGPCR for precise presynaptic inhibition that expands and complements the collection of established and recently developed presynaptic optogenetic tools^16,57^. For manipulating the presynapse, *Pd*CO could potentially serve as a suitable template to create optoGPCR chimeras with altered signaling specificity by exchanging the intracellular GPCR interface as previously demonstrated for other rhodopsin GPCRs^58–68^. *Pd*CO’s biophysical properties are highly suitable for one-photon spectral multiplexing approaches, which are becoming more common in the systems neuroscience field. We believe that *Pd*CO, along with existing optogenetic sensors and with future improved, red-shifted indicators of neuronal activity, will serve as a valuable tool that will allow a better understanding of long-range neural communication in the brain.

## Acknowledgments

We thank all of our lab members and collaborators for thoughtful discussions and comments on the manuscript. This work is supported by grants from the Deutsche Forschungsgemeinschaft (DFG; German Research Foundation) SPP 1926 (B.R.R, O.Y., J.S.W, P.S.), EXC-2049 – 390688087, SPP 1665, SFB 958 (to D.S.), SFB 1315 (to B.R.R and D.S.), the National Institutes of Health (1U01NS128537-01 to B.C. and O.Y.), the European Horizon 2020 Program (ERC CoG PrefrontalMap 819496 and H2020-RIA DEEPER 101016787 to O.Y., ERC SyG BrainPlay 810580 to D.S., ERC-2016-StG 714762 to J.S.W., ERC-2019-CoG 864353 to Y.N.), and the Israel Science Foundation (ISF 3131/20 to O.Y, ISF 1557/2 to Y.N.). J.W. is supported by the EMBO ALTF 378-2019 and Amos de Shalit-Minerva fellowship. O.Y. is supported by the Joseph and Wolf Lebovic Charitable Foundation Chair for Research in Neuroscience. J.D. is the incumbent of the Achar research fellow chair in electrophysiology. We thank Asuka Inoue (Tohoku University) for delta-G7 HEK293 cells, Emilia Entcheva, Harald Janoviak, Mudi Sheves, Moran Shalev-Benami, Stefan Herlitze, Peter Hegemann, Sebastian Bitzenhofer, Johannes Vierock, and Rob Lucas for fruitful discussions and sharing further materials.

## Online methods

### Animals

All experiments involving animals were carried out according to the guidelines stated in directive 2010/63/EU of the European Parliament on the protection of animals used for scientific purposes. Animal experiments at the Weizmann Institute of Science were approved by the Institutional Animal Care and Use Committee (IACUC) of the Weizmann Institute; experiments in Berlin were approved by the Berlin local authorities and the animal welfare committee of the Charité-Universitätsmedizin Berlin, Germany. Experiments in Bonn and Hamburg were performed in accordance with the guidelines of local authorities. Experiments in Tel Aviv were approved by the Institutional Animal Care and Use Committee (IACUC) of Tel Aviv University (approval 01-19-037). For the locomotor behavior experiments, male and female mice (DAT-IRES-Cre; The Jackson Laboratory, Strain #006660) were used. Mice were housed in groups, 2-5 littermates of the same sex per cage. Cagemates underwent surgery on the same day and were assigned to the *Pd*CO or control group such that cages always included mixed groups. The control group included 13 mice (10 males and 3 females, 8-27 weeks old at the time of surgery). The *Pd*CO group included 14 mice (10 males and 4 females, 10-27 weeks old at the time of surgery). For in vivo pupillometry experiments, male and female DBH-Cre (B6.FVB(Cg)-Tg(Dbh-cre)KH212Gsat/Mmucd; 036778-UCD-HEMI) mice (8-12 weeks old at the time of surgery) were used. All mice (DAT-IRES-Cre and DBH-Cre) were kept at 22±2°C, 55±10 % room humidity in a 12-h light/dark cycle with access to food and water ad libitum. Mice were checked daily by animal caretakers.

### Molecular biology and DNA constructs

Mammalian codon optimized Genes encoding optoGPCRs were synthetized (Twist biosciences, USA; except for *Lc*PPO which was generously provided by P. Hegemann, Humboldt-Universität zu Berlin) and fused to a C-terminal rhodopsin 1D4 tag (TETSQVAPA). All genes were further subcloned in-frame with a C-terminal mScarlet into pcDNA3.1 vector under a CMV promoter or into pAAV vector under the CaMKIIα minimal promotor (CaMKIIα 0.4). Expression vectors additionally contained the Kir2.1 membrane trafficking signal (KSRITSEGEYIPLDQIDINV) and Kir2.1 ER export signal (FCYENEV)^69^, N- and C-terminal to mScarlet, respectively, as previously reported for *As*OPN3^13^. The following genes were used for expression (NCBI GeneBank identifier; modifications if applied): *Gg*Opn5l1^70^ (AB368181; modified N- and C-termini originating from *Xenopus tropicalis* OPN5m to improve expression as reported elsewhere^28^ and further C-terminally extended by the last 25 amino acids of *As*OPN3), *Bp*OPN^71^ (AB050606.1), *Tr*PPO2^72^ (AB626965), *Ol*TMT1A^73^ (AGK24990; C-terminus truncated by 63 amino acids to improve expression as reported elsewhere^29^), *Gg*OPN5m^70^ (AB368182), *Hs*OPN5m^74^ (AY377391; C-terminus truncated by 37 amino acids to improve expression as reported elsewhere^75^), *Lc*PPO^23^ (BAD13381), *Pd*CO^25^ (AY692353), *Tr*PPO1^72^ (AB626964), *As*OPN3^22^ (BAN05625; C-terminus truncated by 99 amino acids to improve expression as reported elsewhere ^13,22^) and *Dr*PPO1^72^ (AB626966). CAG-FR-GECO1c was a gift from Robert Campbell (Addgene plasmid #163682)^14^, subcloned into pAAV vector under CaMKIIα 0.4 promotor. The pcDNA3.1-CMV-GIRK2.1 plasmid was a gift from Eitan Reuveny, Weizmann Institute of Science. Gsx^27^ and TRUPATH^30^ plasmids were obtained from Addgene (Gsx: #109373, #109375, #109373, #109360, #109359, #109357, #109356, #109355, and #109350, TRUPATH Kit # 1000000163). Further subcloning into other expression vectors and substitution of mScarlet by EGFP were performed by PCR and/or restriction enzyme-based cloning or the Gibson assembly method^76^. For rAAV packaging limitations the standard woodchuck hepatitis posttranscriptional regulatory element (WPRE) and bovine growth hormone polyadenylation signal (bGHpA) were replaced by a size-optimized expression cassette (miniWPRE, CW3SL)^77^.

The following constructs are available from Addgene(#ID): pcDNA3.1-CMV-PdCO-mScarlet (#198507), pAAV-CaMKIIa(0.4)-PdCO-mScarlet-WPRE (#198508), pAAV-hSyn-SIO-PdCO-mScarlet-WPRE (#198509), pAAV_hSyn-PdCO-mScarlet-WPRE (#198510), pAAV_hSyn-DIO-PdCO-mScarlet-WPRE (#198511), pcDNA3.1-CMV-PdCO-EGFP (#198512), pAAV_hSyn-PdCO-EGFP-WPRE (#198513), pAAV_CaMKIIa(0.4)-PdCO-EGFP-WPRE (#198514), pAAV_EF1a-fDIO-PdCO-mScarlet-miniWPRE (#198515), pAAV_EF1a-DIO-PdCO-mScarlet-ER-miniWPRE (#198516), pAAV-CaMKIIa(0.4)-stChrimsonR-EGFP-P2A-PdCO-WPRE (#202198), and pAAV_hSyn-SIO-stChrimsonR-EGFP-P2A-PdCO-miniWPRE (#202199; all available at https://www.addgene.org/Ofer_Yizhar/). Other constructs are available from the authors upon request.

### Recombinant AAV vector production

For production of rAAV particles, human embryonic kidney cells (HEK293T) cells were seeded at 30±5% confluency and transfected 1d post seeding with plasmids encoding for AAV rep, caps of AAV2 and AAV1 or AAV9 and a vector plasmid encoding for rAAV cassette expressing the above-described optoGPCRs using the PEI method^78^. 72h post transfection, cells were harvested and concentrated by centrifugation at 300g. The resulting cell pellet was resuspended in lysis solution (in mM): 150 NaCl, 50 Tris-HCl; pH 8.5 with NaOH). Cell lysis was performed by three freeze-thaw cycles and treated with 250 U/ml lysate benzonase (Sigma) at 37°C for 1.5h to remove genomic and unpacked DNA, followed by centrifugation at 3000 g for 15 min. Crude virus used for transducing neuronal cultures was filtered with sterile 0.45 µm PVDF filters (Millex-HV, Merck). To produce purified rAAVs, the virus-containing supernatant (crude rAAV) was purified using heparin-agarose columns, eluted with 0.5M NaCl and washed with phosphate buffered saline (PBS). The resulting viral suspension was concentrated with 100 kDa Ultra-15 centrifugal filters (Amicon), aliquoted, and stored at -80°C. Viral titers were quantified by real-time PCR.

### G-protein coupling assays using human embryonic kidney cell culture

For initial testing of Gα signaling specificity, optoGPCR variants expressed in HEK293ΔG7 (lacking GNAS/GNAL/GNAQ/GNA11/GNA12/GNA13/GNAZ; A. Inoue, Tohoku University, Japan)^79^ were tested using the GsX live cell assay^27^. In brief, cells were grown at 37°C, 5% CO_2_ in DMEM containing 4500 mg/L glucose, L-glutamine (Sigma Aldrich) with penicillin/streptomycin (100 U/ml) and 10% FBS. Cells were seeded (2.5 x 10^4^ cells/well) into poly-L-Lysine (Sigma Aldrich) coated solid white 96-well plates (Greiner) and were co-transfected with different opto-GPCR variants (pcDNA3.1, 50 ng/well) together with individual G-protein chimera (GsX, 2 ng/well) and Glo22F Luciferase (GloSensor, Promega, 100 ng/well) using Lipofectamine 2000 (ThermoFisher). Cells were incubated for 24 h at 37°C, 5% CO_2_ and subsequently, in Leibovitz’s L-15 media (without phenol-red, with L-glutamine, 1% FBS, penicillin/streptomycin 100 mg/ml), 9-cis retinal (10 µM) and beetle luciferin (2 mM in 10 mM HEPES pH 6.9) for 1 h at RT. Cells were kept dark and baseline luminescence was measured over a time period of 200 s followed by opto-GPCR activation using a 1 s light pulse (collimated CoolLED pE4000, Andover, UK) of either 385 nm or 470 nm (for *Tr*PPO2, *Ol*TMT1A, and *Bb*OPN). Changes in cAMP levels were measured over time using GloSensor luminescence with a Mithras Luminometer (Berthold Technologies, Germany). For the assay quantification each biological replicate was normalized to its pre-light baseline as well as to a non-optoGPCR control.

For the testing of optoGPCR-mediated inhibition of adenylate cyclase activity HEK293ΔG7 cells were seeded and transfected as described above with (plasmid amount per well in ng: 100 GloSensor, 2 Gαs, 50 optoGPCR). 24 h after transfection, the medium was changed to phosphate-buffered saline (with Ca2+ and Mg2+), supplemented with 9-cis retinal (10 µM) and beetle luciferin (2 mM in 10 mM HEPES pH 6.9). After 1 h incubation at RT, the baseline bioluminescence was measured, followed by application of forskolin (500 µM final concentration). After 20 min, optoGPCRs were activated with a 2 s light pulse (365 nm) and bioluminescence measurements were continued for another 60 mins. Bioluminescence signals were normalized to the average bioluminescence prior forskolin application. For more detailed Gα-protein profiling, the TRUPATH assay^30^ was used and HEK293ΔG_7_ cells were seeded as described above, co-transfected with RLuc8-Gα, Gβ, Gγ-GFP2 and optoGPCRs in a 1:1:1:1 ratio (100 ng/well total DNA) using Lipofectamine 2000. Cells were incubated for 24 h at 37°C, 5% CO_2_ and subsequently, in Leibovitz’s L-15 media (without phenol-red, with L-glutamine, 1% FBS, penicillin/streptomycin 100 mg/ml) and 9-cis retinal (10 µM) and kept in the dark. For performing BRET assays the medium was changed to HBBS, supplemented with 20 mM HEPES, 10 μM 9-cis-retinal + 5 μM Coelenterazine 400a, and incubated for 5 min at RT. optoGPCRs were activated using a 1 s, 385 nm light pulse (collimated CoolLED pE4000, Andover, UK). BRET ratio changes were determined from RLuc8-Gα and Gγ-GFP2 signals, integrated over 3 minutes, directly after light application and 17 to 20 post optoGPCR activation.

### optoGPCR mediated GIRK current recordings from human embryonic kidney cells

For the initial comparison of optoGPCR-evoked GIRK currents, optoGPCRs were transiently expressed in HEK293 cells stably expressing GIRK1/2 subunits (kindly provided by Dr. A. Tinker, Queen Mary’s School of Medicine and Dentistry). Briefly, cells were maintained at 37°C, 5% CO_2_ in high glucose DMEM supplemented with Geniticin (G418, GIBCO), 10% fetal bovine serum (FBS, Biological Industries) and penicillin/streptomycin (100 U/ml) and seeded onto poly-D-lysine coated coverslips in 24-well plates (Corning) and were additionally supplemented with 1µM 9-cis retinal (Sigma). One day post-seeding, pcDNA3.1-CMV-optoGPCR-mScarlet plasmids were transiently transfected using FuGENE HD (Promega; 0.75 µl/well, plasmid DNA, 250 ng/well) in serum free DMEM (50 µl/well).

Currents from HEK293 cells stably expressing GIRK were recorded under visual guidance using a slice scope II (Scientifica, UK) with an Olympus LUMPlanFL N 40x/0.80 W objective under IR-DIC. A Lumencor SpectraX light engine was used to identify expressing cells via mScarlet fluorescence and for light application to toggle optoGPCR activation. In the case of non-switchable or slow cycling optoGPCRs (*As*OPN3, *Ol*TMT1A, *Gg*OPN5l1), expressing cells were identified first and patched only after an additional 25 min in darkness. HEK cells were perfused with extracellular solution (in mM): 20 NaCl, 120 KCl, 2 CaCl_2_, 1 MgCl_2_, 10 HEPES, pH 7.3 (KOH), 320 mOsm (with D-glucose). Glass microelectrodes (1.5-2.5 MΩ) were pulled from thin-walled glass capillaries and filled with (in mM): 5 NaCl, 40 KCl, 2 MgCl_2_, 10 HEPES, 100 K-aspartate, 5 MgATP, 0.1 Na_2_GTP, 2 EGTA, pH 7.3 (KOH), 300 mOsm (with D-glucose). GIRK currents were recorded in whole-cell voltage-clamp mode at a holding potential of -50 mV. A Multiclamp 700B amplifier and Digidata 1440A digitizer (both Molecular Devices) were used to control and acquire electrophysiological recordings. Data was acquired at 10 kHz and filtered at 3 kHz.

The different optoGPCRs were activated with a 500 ms (5 s in case of *Gg*OPN5l1) light pulse close to their reported activation maximum with 10 nm narrow bandpass filters (Edmund optics). Light intensities for each wavelength were calibrated to the same photon flux corresponding to 0.92 mW/mm² at 520nm. The following center wavelengths were used (in nm): 520 *As*OPN3, 450 *Ol*TMT1A, 473 *Tr*PPO2, 450 *Bb*OPN. 546 *Gg*OPN5l1, and 394 for all other optoGPCRs. Light induced recovery was induced by application of a 10 s 568±10 nm light pulse. Experiments were performed at 22±1°C. Maximum GIRK current amplitudes were determined using Clampfit 10.7 (Molecular Devices). Light intensities were measured with a calibrated S170C power sensor (Thorlabs).

For two-photon activation of *Pd*CO, electrophysiological recordings were performed on HEK293T cells (HEK293T/17, ATCC, #CRL-1573) as described previously^14^. In brief, pcDNA3.1-CMV-PdCO-mScarlet was co-transfected (1:3) together with pCAG-GIRK2/1-myc^80^ using Lipofectamine 2000 (Invitrogen) according to manufacturer instructions and supplemented with 1 µM 9-cis retinal (Sigma). GIRK currents evoked by activation of PdCO were recorded under visual guidance using a Fluoview FVMPE-RS multiphoton imaging system using an XLPLN25XWMP2 objective (both Olympus). The extracellular solution contained (in mM): 140 NaCl, 20 KCl, 0.5 CaCl_2_, 2 MgCl_2_, 10 Glucose, 10 HEPES, pH to 7.3 with NaOH, 315±5 mOsm. Cells were patched with microelectrodes pulled from thin-walled glass capillaries (1-5 MΩ) and filled with (in mM): 120 K-gluconate, 5 NaCl, 0.1 CaCl_2_, 2 MgCl_2_, 1.1 EGTA, 10 HEPES, 4 Na_2_ATP, 0.4 Na_2_GTP, 15 Na_2_-phosphocreatine, pH=7.28, 290 mOsm. Whole-cell voltage-clamp current recordings in response to 200 ms ramp depolarizations from -120 to 40mV every 2s with a holding potential of -40mV were amplified and digitized (20 kHz) using a HEKA EPC10 (filtered at 3 kHz). Whole-cell and pipette capacitance transients were minimized, and series resistance was compensated by 70%.

Two-photon excitation for *Pd*CO was carried out using MaiTai and Insight tunable Ti:Sapphire lasers (Spectra Physics). For photostimulation, cells were centered in a 90×90 pixel square (0.4792mm/pixel) and scanned at a speed of 8µs/pixel. The spectral characterization was performed using a 20 s two-photon stimulation at 700, 800, 900, 1000, or 1100nm at 3mW laser power. For each cell, 3 to 5 different wavelengths were applied randomly. For a dose-response titration at 800 nm, a 10 s stimulation was performed across 0.1, 0.3, 1, 3, and 10 mW from low to high intensity. Two-photon intensities were calibrated at the sample focal plane using a thermal power sensor (S175C, Thorlabs) and power meter (PM100D, Thorlabs). Data was analyzed using IgorPro (Wavemetrics) and NeuroMatic^81^ using custom macros. For each voltage ramp, the GIRK-mediated inward current was averaged over 5 ms at -120 mV holding potential.

### Primary dissociated hippocampal neuron culture and gene delivery

Primary cultured hippocampal neurons were isolated from CA1 and CA3 hippocampal regions of either sex P0 Sprague-Dawley rat pups (Envigo). Neurons were digested with 0.4 mg/ml papain (Worthington) and seeded on Matrigel (1:30, Corning) coated glass coverslips in 24-well plates at a density of 65,000 cells per well. Neurons were maintained in a 5% CO_2_ humidified incubator in Neurobasal-A medium (Invitrogen) supplemented with 1.25% FBS, 4% B-27 supplement (GIBCO), and 2 mM Glutamax (GIBCO). For inhibition of glial overgrowth, 200 mM fluorodeoxyuridine (Sigma) was added at DIV 4 (days in vitro).

For confocal imaging or initial electrophysiological recordings of opsin expressing cultured primary neurons, opsin and cell-filling EYFP or GIRK2.1 encoding plasmids were co-transfected at DIV5 using a modified Ca^2+^-phosphate method^82^. Briefly, the neuronal cultured medium of a 24 well plate was collected and replaced with 400 ml serum-free modified eagle medium (MEM, Thermo Fisher Scientific). 30 µl transfection mix (2 mg plasmid DNA and 250 mM CaCl_2_ in HBS at pH 7.05) was added per well, and cells were incubated for 1h to allow for transfection. Neurons were washed twice with MEM, and the medium was changed back to the collected original medium. Cultured neurons were used between 14-17 DIV for experiments.

For Ca^2+^ imaging experiments, cultured neurons were co-transduced with pAAV_CaMKIIα(0.4)-PdCO-EGFP-WPRE and pAAV_CaMKIIα(0.4)-FR-GECO1c-WPRE at DIV 1. Experiments were carried out between 14-21 DIV.

Autaptic primary hippocampal neuronal cultures on glia cell micro-islands were prepared from P0 mice (C57BL/6NHsd; Envigo, Cat#044) of either sex as previously described^83^. 300 µm diameter spots of growth permissive substrate consisting of 0.7 mg ml^-1^ collagen and 0.1 mg ml^-1^ poly-D-lysine were applied with a custom-made stamp on agarose-coated coverslips. First, astrocytes were seeded and were allowed to proliferate in Dulbecco’s modified eagle medium (DMEM) supplemented with 10% fetal calf serum and 0.2% penicillin/streptomycin (Invitrogen) for one week to form glia micro-islands. After changing the medium to Neurobasal-A supplemented with 2% B27 and 0.2% penicillin/streptomycin, hippocampal neurons prepared from P0 mice were added at a density of 370 cells cm^-2^. Neurons were transduced with rAAVs (1.5 10^8^ VG per well, titer-matched for all constructs) at DIV 1 and were recorded between DIV 14 and DIV 21.

### Hippocampal organotypic slice culture and gene delivery

Organotypic hippocampal slices were prepared from Wistar rats at postnatal day 5-7 as described^84^. Briefly, dissected hippocampi were cut into 400 μm slices with a tissue chopper and placed on a porous membrane (Millicell CM, Millipore). Cultures were maintained at 37°C, 5% CO_2_ in a medium containing 80% MEM (Sigma M7278), 20% heat-inactivated horse serum (Sigma H1138) supplemented with 1 mM L-glutamine, 0.00125% ascorbic acid, 0.01 mg/ml insulin, 1.44 mM CaCl_2_, 2 mM MgSO_4_ and 13 mM D-glucose. No antibiotics were added to the culture medium.

Transgene delivery in individual CA3 pyramidal cells was performed by single-cell electroporation between DIV 15-20 as previously described^85^. The plasmids pAAV-CaMKIIα(0.4)-PdCO-mScarlet and pAAV-CaMKIIα(0.4)-LcPPO-mScarlet were each diluted to 50 ng/ml in K-gluconate-based solution consisting of (in mM): 135 K-gluconate, 10 HEPES, 0.2 EGTA, 4 Na_2_-ATP, 0.4 Na-GTP, 4 MgCl_2_, 3 ascorbate, 10 Na_2_-phosphocreatine, pH 7.2, 295 mOsm/kg. An Axoporator 800A (Molecular Devices) was used to deliver 25 hyperpolarizing pulses (-12 V, 0.5 ms) at 50 Hz. During electroporation, slices were maintained in pre-warmed (37°C) HEPES-buffered solution consisting of (in mM): 145 NaCl, 10 HEPES, 25 D-glucose, 2.5 KCl, 1 MgCl_2_ and 2 CaCl_2_ (pH 7.4, sterile filtered). Targeted viral vector-based transduction of organotypic hippocampal slice cultures^86^ was performed by pressure injecting (20 PSI/2-2.5 bar, 50 ms duration) recombinant adeno-associated viral particles encoding AAV2/9.CaMKIIα(0.4)-PdCO-mScarlet or AAV2/9.CaMKIIα(0.4)-LcPPO-mScarlet and AAV2/9.hSyn-somBiPOLES-mCerulean using a Picospritzer III (Parker) under visual control (oblique illumination) into CA3 *stratum pyramidale* between DIV 2-5. Slice cultures were then maintained in the incubator for 2-3 weeks allowing for virus payload expression.

### Confocal imaging, quantification membrane targeting and expression levels

Primary cultured hippocampal neurons (transfected as described above) co-expressing the different opsins (pAAV-CaMKIIα(0.4)-opsin-mScarlet) together with a cell filling EYFP (pAAV-CaMKIIα(0.4)-EYFP) were fixed and permeabilized 4 days post transfection using 4% PFA for 15 min, washed 3 times with PBS and stained for 3 min with DAPI (5 mg/mL solution diluted 1:30,000). Coverslips were mounted using PVA-DABCO (Sigma) and fluorescence images were acquired using a Zeiss LSM 700 confocal microscope equipped with a Plan-Apochromat 63x/1.40 Oil DIC objective.

For quantification of opsin expression in the membrane and cytosol, respectively, binary masks for EYFP and mScarlet signals were generated using fixed thresholding in imageJ^87^ on a single equatorial z-slice per expressing neuron, identified visually with help of the nuclear DAPI stain. Expression analysis was restricted to the somatodendritic region by manual selection. The EYFP mask was subtracted from the mScarlet mask to generate a mask that restricts the analysis to the membrane only. Subsequently, the average pixel intensity was measured for the defined regions of interest. The expression index was calculated by subtraction of the whole cell mScarlet signal by the EYFP signal, divided by the sum of both signals.

### In vitro electrophysiology on neuronal samples

Qualitative measurements of optoGPCR functionality using primary cultured neurons co-expressing pAAV-CaMKIIα(0.4)-optoGPCR-mScarlet together with pcDNA3.1-CMV-GIRK2.1 transfected as described above, were performed using the same setup as described for measurements on stably expressing GIRK2/1 HEK293 cells. Neurons were patched with microelectrodes (3.0-4.5 mΩ), filled with (in mM): 2 NaCl, 4 KCl, 10 HEPES, 135 K-gluconate, 4 Na_2_ATP, 4 EGTA, 0.3 Na_2_GTP, 290 mOsm, pH adjusted to 7.3 with KOH. Electrophysiological recordings obtained under continuous perfusion in Tyrode’s medium (in mM): 150 NaCl, 4 KCl, 2 MgCl_2_, 2 CaCl_2_, 10 D-glucose, 10 HEPES; 320 mOsm; pH adjusted to 7.35 with NaOH.

EPSCs from autaptic primary neurons were recorded under visual guidance using an Olympus IX51 inverted microscope using an Olympus UPlanSApo 20x/0.75 UIS2 objective under far infrared light (>665 nm) widefield illumination. A CoolLED P4000 served as a light source to identify expressing cells and for light application to toggle optoGPCR activation. In the case of non-switchable optoGPCRs (*As*OPN3 and *Ol*TMT1A), electrophysiological recordings were performed first, and cells were investigated for expression post recordings. Acquired data was excluded in case cells were not expressing. Autaptic neurons were constantly perfused with extracellular solution (in mM): 140 NaCl, 2.4 KCl, 10 HEPES, 10 D-glucose, 2 CaCl_2_, and 4 MgCl_2_ (pH adjusted to 7.3 with NaOH, 300 mOsm). Cells were patched with microelectrodes pulled from quartz glass capillaries (3-4 mΩ), filled with (in mM): 136 KCl, 17.8 HEPES, 1 EGTA, 0.6 MgCl_2_, 4 MgATP, 0.3 Na_2_GTP, 12 Na_2_ phosphocreatine, 50 U/ml phosphocreatine kinase (300 mOsm); pH adjusted to 7.3 with KOH. A Multiclamp 700B (Molecular Devices) amplifier and NI USB-6343 digitizer (National Instruments) were used to control and acquire electrophysiological recordings and the application of light stimulation via WinWCP software (University of Strathclyde, Glasgow). Data was acquired at 10 kHz and filtered at 3 kHz. Cells were kept at -70 mV and series resistance and capacitance were compensated by 70%. EPSCs were elicited by a 1 ms depolarization to 0 mV (50 ms interstimulus interval, every 5 s) resulting in an unclamped axonal action potential causing neurotransmitter release. SCH23390 (Tocris) was locally applied with a perfusion system (AutoMate Scientific ValveLink8.2). Pertussis toxin (0.5 mg/ml) was applied to the cultures 24 h before the recordings. For the initial comparison of EPSC reduction efficacy, the different optoGPCRs were activated with a 0.5 s, 390±10 nm (FB390-10, Thorlabs) and potential recovery was induced by a 4.5 s, 560±10 nm (FB560-10, Thorlabs). For spectral sensitivity measurements, light from the CoolLED P4000 was filtered with narrow bandpass filters mounted on a FW212C filter wheel (Thorlabs). The following filters were used (center wavelength ±10 nm, [Edmund optics catalog no.]): 365 [#65-069], 405 [#65-072], 436 [#65-077], 470 [#65-083], 492 [#65-087], 520 [#65-093], 560 [#87-887], 594 [#86-733], 636 [#65-106], and 660 [#86-086]. Light intensities for each wavelength were calibrated to the same photon flux corresponding to 1.1 mW/mm² at 520nm. Light intensities were measured with a calibrated S130VC power sensor (Thorlabs). Spectral measurements were performed alternating from UV to red wavelengths or vice versa. Light titration experiments were performed from low to high light intensity or light pulse duration. Experiments were performed at room temperature.

Electrophysiological recordings in organotypic hippocampal slice cultures were performed using a BX 51WI microscope (Olympus) equipped with a Multiclamp 700B amplifier (Molecular Devices) controlled by Ephus^88^. Alternatively, a second BX 51WI microscope (Olympus) equipped with a Double IPA integrated patch amplifier controlled by SutterPatch software (Sutter Instrument) was used for electrophysiological measurements. Patch pipettes with a tip resistance of 3-5 MΩ were filled with (in mM): 135 K-gluconate, 4 MgCl_2_, 4 Na_2_-ATP, 0.4 Na-GTP, 10 Na_2_-phosphocreatine, 3 ascorbate, 0.2 EGTA, and 10 HEPES (pH 7.2). Artificial cerebrospinal fluid (ACSF) consisted of (in mM): 135 NaCl, 2.5 KCl, 4 CaCl_2_, 4 MgCl_2_, 10 Na-HEPES, 12.5 D-glucose, 1.25 NaH_2_PO_4_ (pH 7.4). All experiments were performed at room temperature (21-23°C) except the extracellular field stimulation experiments, performed at 33±1 °C. For experiments measuring GIRK currents, synaptic blockers D-CPP-ene (10μM), NBQX (10μM), Picrotoxin (100μM) (Tocris, Bristol, UK) were added to the HEPES-buffered ASCF and patched optoGPCR-expressing CA3 neurons were held at -70 mV during the measurements. In synaptic stimulation experiments (optogenetic and electrical), postsynaptic non-transfected CA1 neurons were held at -60 mV while recording PSCs in voltage clamp mode. Access resistance of the recorded CA1 neuron was continuously monitored and recordings above 20 MΩ and/or with a drift > 30% were discarded. A 16-channel pE-4000 LED light engine (CoolLED, Andover, UK) was used for optogenetic stimulation of the optoGPCRs. Light intensity was measured in the object plane with a 1918 R power meter equipped with a calibrated 818 ST2 UV/D detector (Newport, Irvine CA) and divided by the illuminated field of the Olympus LUMPLFLN 60XW objective (0.134 mm2) or of the Olympus LUMPLFLN 40XW objective (0.322 mm^2^). For presynaptic somBIPOLES stimulation we used a fiber-coupled LED (400 µm fiber, NA 0.39, M118L02, ThorLabs) to deliver 5 ms red light pulses at 625 nm.

In extracellular electrical stimulation experiments, afferent Schaffer collateral axons were stimulated (0.2 ms, 20-70 µA every 10 s) with a monopolar glass electrode connected to a stimulus isolator (IS4 stimulator, Scientific Devices).

### One-photon spectral multiplexing in cultured neurons

For calcium imaging experiments paired with transient *Pd*CO activation, hippocampal neuronal cultures were co-transduced with pAAV-hSyn1(0.4)-PdCO-EGFP and pAAV-CaMKIIα(0.4)-FR-GECO1c as described above and experiments were carried out between DIV 14-21. Imaging was performed with SliceScope II (Scientifica, UK) using an Olympus LUMPlanFL N 40x/0.80 W objective and data were acquired with an ORCA-Flash4.0 digital camera (C11440, Hamamatsu). Data acquisition was carried out with 4×4 binning, 80 ms exposure at 10 Hz and FR-GECO1c was excited at 586±10 nm with an irradiance of 0.52 mW/mm². To transiently activate *Pd*CO, 445±10 nm light (6.37 mW/mm²) was applied at 10 Hz, 5 ms pulse width between the imaging intervals. Cells were constantly perfused with extracellular solution (in mM): 140 NaCl, 2.4 KCl, 10 HEPES, 10 D-glucose, 2 CaCl_2_, and 4 MgCl_2_ (pH adjusted to 7.3 with NaOH, 300 mOsm). To evoke AP-induced calcium transients, cells were patched in the tight-seal, cell-attached configuration as described by Perkins^89^. Briefly, cells were patched with low resistance microelectrodes pulled from glass capillaries (1.5-2.5 mΩ), filled with 145 mM NaCl. A train of APs was evoked by a 40 ms square current injection of 0.6 to 1.0 µA at a frequency of 0.2 Hz through the intact membrane patch (>3 GΩ seal resistance). The change in the FR-GECO1c signal ΔF/F_0_ (where F_0_ is the median fluorescence signal and ΔF=F(t)-F_0_) was calculated in imageJ^87^ and corrected for blue light induced increase in the fluorescence signal (Extended Data Fig. 10). A Lumencor SpectraX light engine served as light source and light intensities were measured with a calibrated S170C power sensor (Thorlabs). The spectra of light used to excite *Pd*CO and FR-GECO1c were measured with an Ocean QE pro spectrometer (Ocean Optics).

Experiments using pAAV-CaMKIIα(0.4)-stChrimsonR-EGFP-P2A-PdCO-WPRE were performed on the setup described above using the same extracellular solution. Neurons were patched at -70mV with microelectrodes (3.0-4.5 mΩ), filled with (in mM): 2 NaCl, 4 KCl, 10 HEPES, 135 K-gluconate, 4 Na_2_ATP, 4 EGTA, 0.3 Na_2_GTP, 290 mOsm, pH adjusted to 7.3 with KOH. For action spectra recordings, light from the Lumencor SpectraX light engine was filtered with narrow bandpass filters mounted on a FW212C filter wheel (Thorlabs). The following filters were used (center wavelength ±10 nm, [Edmund optics catalogue no.]): 394 [#65-070], 422 [#34-496], 450 [#65-079], 473 [#34-502], 500 [#65-088], 520 [#65-093], 546 [#65-097], 568 [#65-099], 594 [#86-733], 620 [#65-104], and 647 [#65-108]. The light intensity for each wavelength was calibrated to the same photon flux corresponding to 0.92 mW/mm² at 520nm. Short light pulses were applied (2 ms) and photocurrents measured under TTX (1µM), CNQX (20µM), AP-5 (50µM), recorded in both directions per expressing cell (red to blue and blue to red) and then averaged. For postsynaptic recording in non-expressing cells, neurons were measured with 10µM SCH233900 (10µM) to eliminate the potential contribution of GIRK currents. The light from the Lumencor SpectraX light engine was filtered with bandpass filters and the following light intensities were used: 390±9nm (0.155 mW/mm^2^) 512±12nm (0.327 mW/mm^2^) 632±11nm (7.015 mW/mm^2^). A Multiclamp 700B amplifier and Digidata 1440A digitizer (both Molecular Devices) were used to control and acquire electrophysiological recordings. Data was acquired at 10 kHz and filtered at 3 kHz. Light intensities were measured with a calibrated S170C power sensor (Thorlabs).

### Neonatal cardiomyocytes and beating frequency measurements

Neonatal mouse atrial cardiomyocytes were dissociated from 1-3 day-old CD1 mouse hearts using the Neonatal Heart Dissociation Kit (Miltenyi Biotec) according to the manufacturer’s instructions. Cardiomyocytes in suspension (10^6^ cells) were transfected by nucleofection (4D-Nucleofector X-Unit, CM-120-150 program, P3 buffer, Lonza) with 5 µg plasmid DNA (pcDNA3.1-CMV-PdCO-mScarlet) and plated on fibronectin coated 24 well plates (µ-Plate ibidi). Video microscopy of beating atrial cardiac cells was performed 3-5 days post transfection in a imaging cell chamber with 5 % CO_2_, 80% humidity and 37°C on a Nikon Eclipse Ti2 microscope with a objective S Plan Fl LWD 20x 0.7 NA objective. Changes in spontaneous beating frequency were recorded under IR light using a CMOS camera (Grasshopper3, Teledyne FLIR) and analyzed online using custom designed software (LabView, National Instruments) as described before^90^. Light stimulation was performed with a LedHUB (Omicron-Laserage) equipped with 385 nm LED (attenuated with a 10 % neutral density filter) and a 555 nm LED. Illumination was controlled by a recording system (PowerLab 4/35 and LabChart8 Software, AD Instruments). The 385 nm light intensity was varied logarithmically from 0.27 µW/mm² to a maximum of 132.94 µW/mm² every 25 s with a duration of 500 ms. A 10 s 555 nm light pulse was applied before the 385 nm pulse and repeated every 15 s. Light intensities were measured with an S130A power meter sensor at the objective level (Thorlabs).

### Surgeries

DAT-Cre transgenic mice (n=27) were anesthetized (with isoflurane 2.5% by volume for induction and 1-2% for maintenance) and placed in a stereotaxic frame. Surgery was performed under aseptic conditions. rAAV vectors encoding a Cre-dependent *Pd*CO-mScarlet transgene (rAAV2/1&2-hSyn-SIO-PdCO-mScarlet-WPRE; 3.8E12 viral genomes / ml) or eYFP (rAAV2/1&2.EF1a.DIO.eYFP; 2E13 viral genomes / ml) were bilaterally injected into the substantia nigra (Coordinates respective to bregma: AP: - 3.5 mm, ML: + or - 1.4 mm DV: - 4.25 mm; 500 nl per site, at a rate of 100 nl/min). Optical fibers (200 µm diameter, NA 0.5) were bilaterally implanted above the dorsomedial striatum (AP: + 0.6 mm, ML: + or – 1.5 mm DV: - 2.1 mm). C&B MetaBond cement and Dental acrylic were used to fix the fibers to the skull. Mice were allowed to recover for 6-9 weeks to allow for viral expression. Analgesia was administered before the surgery and for at least three days following recovery.

For pupillometry experiments, mice (n=6) were anesthetized (with isoflurane 3% by volume for induction and 1.3% for maintenance) and placed in a stereotaxic frame. Surgery was performed under aseptic conditions. Under microscopic control, a craniotomy was made using a high-speed surgical drill. Two surgeries were performed on each mouse. In the first surgery the virus (rAAV5/2- or 1/2-hSyn-SIO-PdCO-mScarlet-WPRE; 6.1E12 or 3.8E12 viral genomes / ml) was injected to the LC using a UMP3 Microsyringe Injector, a Micro4 Controller pump, and 33-gauge NanoFil needles (World Precision Instruments, USA) unilaterally at a rate of 100 nl/min for 8 min (two injections of 250 nl for a total volume of 500 nl). Coordinates used to target LC: (respective to bregma) anteroposterior (AP): -5.4 mm; medio-lateral (ML): 1 mm; dorso-ventral (DV): -3.3,-3.5 mm relative to brain surface. In the second surgery, optic fibers (MFC_200/240-0.22_10 mm_MF1.25_FLT, Doric Lenses) were implanted above the BF (basal forebrain, AP: 0.7 mm; ML: 2.25 mm at 7.5°, DV: -4.75 mm from brain surface) and above the EW (AP: -3.8 mm; ML: 1.3 mm at 20°; DV: -3.2 mm relative to brain surface). In one animal the CPu (caudate-putamen) was targeted instead of the BF [AP -1.1, ML 1.35, DV -4]. Dental acrylic was gently placed around the optic fibers, fixing them to the skull.

### Pupillometry experiments

Mice were anesthetized with (0.7-1% isoflurane) and placed in a stereotaxic frame. Optic patch cords connected the implanted optic fiber ferrules to a 447 nm laser (Changchun New Industries Optoelectronics Technology Co., Ltd., China). Pupillometry was performed by illuminating the eyes with infrared light (VGAC) and by continuous video monitoring (10 frames/s) synchronized with laser stimulation. We examined the effects of different stimulation frequencies (1,5,10,20 and 40 Hz) in two separate locations (EW or forebrain) on ipsilateral or contralateral pupil size (relative to injection and fiber). Each stimulation trial was 10 seconds long, with 20 ms pulses and a long 2-minute inter-trial interval to allow pupils to return to baseline. Trial order was randomized for stimulation frequency and stimulation location within each experiment. Data analysis (quantification of pupil area in video frames) was performed as previous studies using custom Matlab code^91^. Briefly, we fit a circle to the pupil area in the video images and, for each trial separately, normalized pupil area relative to the 4s pre-trial baseline to compute percent change dynamics in the [−5 30] s interval around laser stimulation. Trials were then averaged for each animal, eye, and condition separately (∼8 trials per condition) to generate time-courses as seen in Fig. 6j. Maximal pupil constriction (through in time-course) was defined as the minimal value in this pupil time-course.

### In vivo optogenetic silencing of the nigrostriatal pathway

Following recovery, mice underwent a single 15-minute habituation session, to habituate to handling, bilateral patch cord attachment and the open field arena. In experimental sessions, we attached individual mice to a patch cord unilaterally to measure *PdCO-induced* bias in locomotion. We recorded the free locomotion in an open field arena (50×50×50 cm) continuously over 30 minutes under near-infrared illumination. After a 10-minute baseline no-light period (“baseline”), we delivered 500 ms light pulses (400 nm, 10 mW at the fiber tip), at 0.4 Hz for 10 minutes (“*Pd*CO activation”), to the right or left sides (on separate sessions). Then we administered 5 light pulses (530 nm, 4 sec duration at 0.1 Hz, 10 mW at the fiber tip; “*Pd*CO recovery”) followed by no-light administration for the rest of the 10 minutes recovery period. A MASTER-8 pulse generator (A.M.P.I.) triggered the activation of both wavelengths, delivered by STSI-Optogenetics-LED-Violet-Green system (Prizmatix). A near-infrared LED placed at the video ROI was used as a synchronization signal (100 ms pulse duration, 0.1 Hz, for the entire 30 minutes session). These digital signals were split and recorded using Open Ephys acquisition board and used to synchronize the video and LED operation.

Offline video processing and mouse tracking was done using DeepLabCut^92^. Briefly, we trained DLC to detect 7 features on the mouse body (nose, head center, left and right ears, left and right hips, tail base), the 4 corners of the arena and the ON or OFF states and coordinates of the video synchronization LED. X-Y coordinates of each feature were then further processed to complete missing or noisy values (large and fast changes in X or Y) using linear interpolation (interp1) of data from neighboring frames. This was followed by a low pass filtering of the signals (malowess, with 50 points span and of linear order). Finally, a pixel to cm conversion was done based on the video-detected arena features and its physical measurements. A linear fit to the nose, head, center and tail features defined the mouse angle with respect to the south arena wall at each frame. Following its dynamics over the session, we identified direction shifts as a direction change in angle that exceeds 20 degrees and 1 second. To achieve a comparable measurement between right- and left-hemisphere sessions, we measured motion in the ipsilateral direction as positive and contralateral motion as negative from the cumulative track of angle. The net angle gain was calculated as the sum of ipsilateral and contralateral angles gained over each time bin (1- or 10-minute bins as indicated). Results from the left- and right hemisphere sessions of each mouse were averaged and then used for statistical comparison between the *Pd*CO and control groups.

### Data analysis, quantification, and statistics

Phylogenetic trees were generated with phylogeny.fr^93^. In vitro electrophysiological recordings in cultured cells were analyzed using Clampfit 10.7 (Molecular Devices) as well as IgorPro (Wavemetrics) and NeuroMatic^81^ for two-photon experiments. Analysis of mEPSCs data was performed using Easy electrophysiology (v2.3.3b) with a 0.37 correlation cutoff and a 15pA amplitude threshold due to artificial noise created by series resistance compensation. G-protein coupling assays (TRUPATH, GsX, GloSensor) were analyzed in Microsoft Excel. Confocal imaging and calcium imaging data was analyzed in ImageJ^87^. Data from organotypic slice recordings was analyzed with MATLAB. Atrial cardiomyocyte beating was analyzed using LabView (National Instruments). In vivo experiments were analyzed using Matlab and DeepLabCut^92^. Statistical analysis was performed with MATLAB, Graphpad Prism 9 and estimation statistics were performed online^94^.

Sample sizes were similar to those commonly used in the field and no statistical tests were run to predetermine sample size. Blinding was performed in autaptic benchmark experiments (Fig. 2e) and behavioral experiments silencing the nigrostriatal pathway. Randomization was performed in the nigrostriatal pathway silencing experiments, for biophysical characterization of optoGPCRs in autaptic neurons, and for two-photon characterization. Automated analysis was used whenever possible. For autaptic neuron recordings, cells were excluded from analysis if the first EPSC amplitude was below 100pA, and from the analysis of the paired-pulse ratio if optoGPCR activation completely abolished the first EPSC. Further, were cells excluded from analysis if the access resistance was above 20 MΩ or if the holding current exceeded 200 pA. For organotypic slice culture recordings cells were additionally excluded from analysis if a EPSC amplitude drift > 30% occurred. For in vivo electrophysiology (Figure 6), recording sessions in which no units showed visual stimulus-evoked activity were excluded from the analysis.

## Extended Data Figures

**Extended Data Fig. 1:**
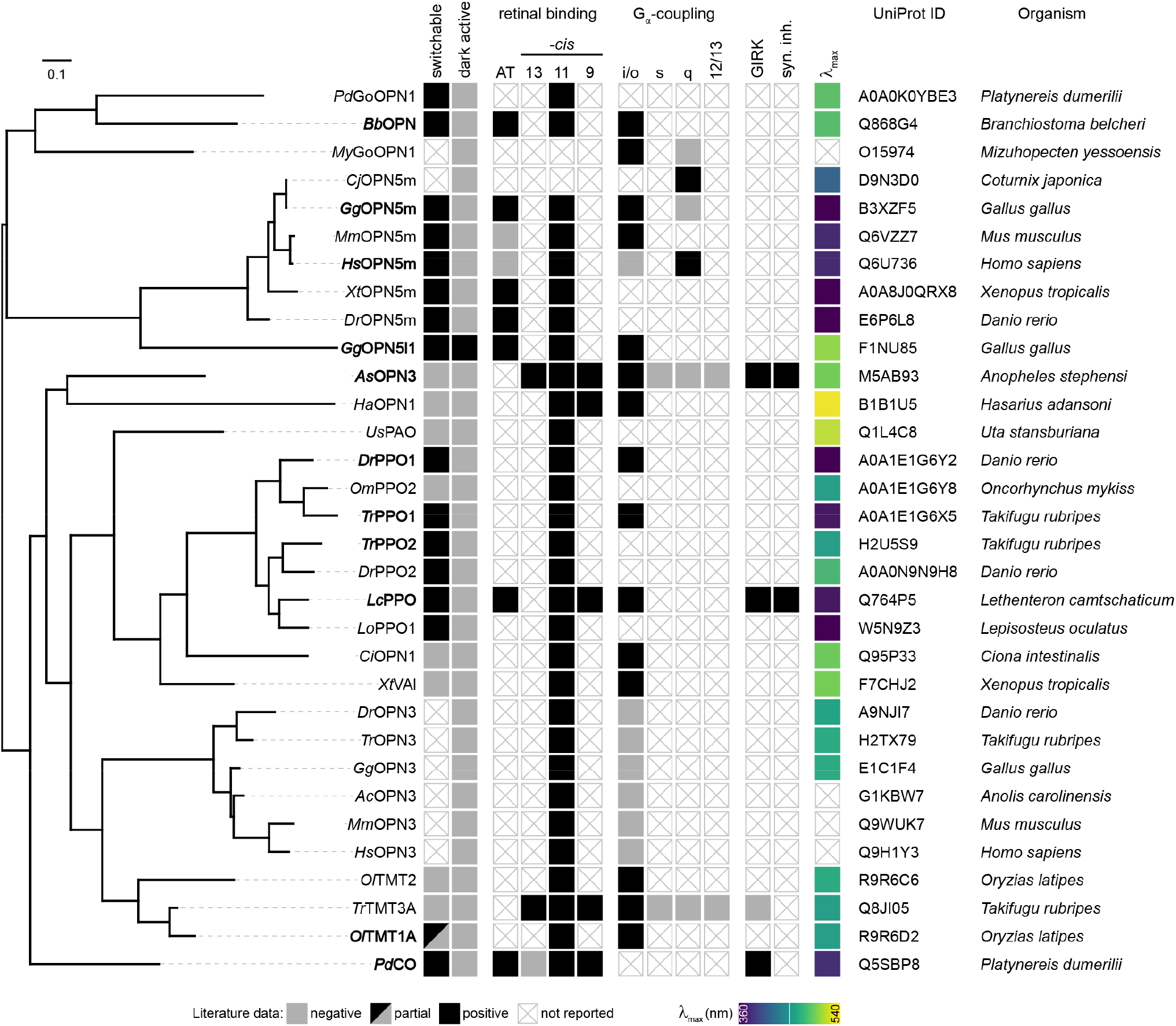
Curated literature data of optoGPCRs. Overview of all optoGPCR and their properties from literature (for references please see source data file). Left: phylogenetic tree. Middle section (squares): optoGPCR properties. Right: UniProt identifiers and species of origin of optoGPCRs.

**Extended Data Fig. 2:**
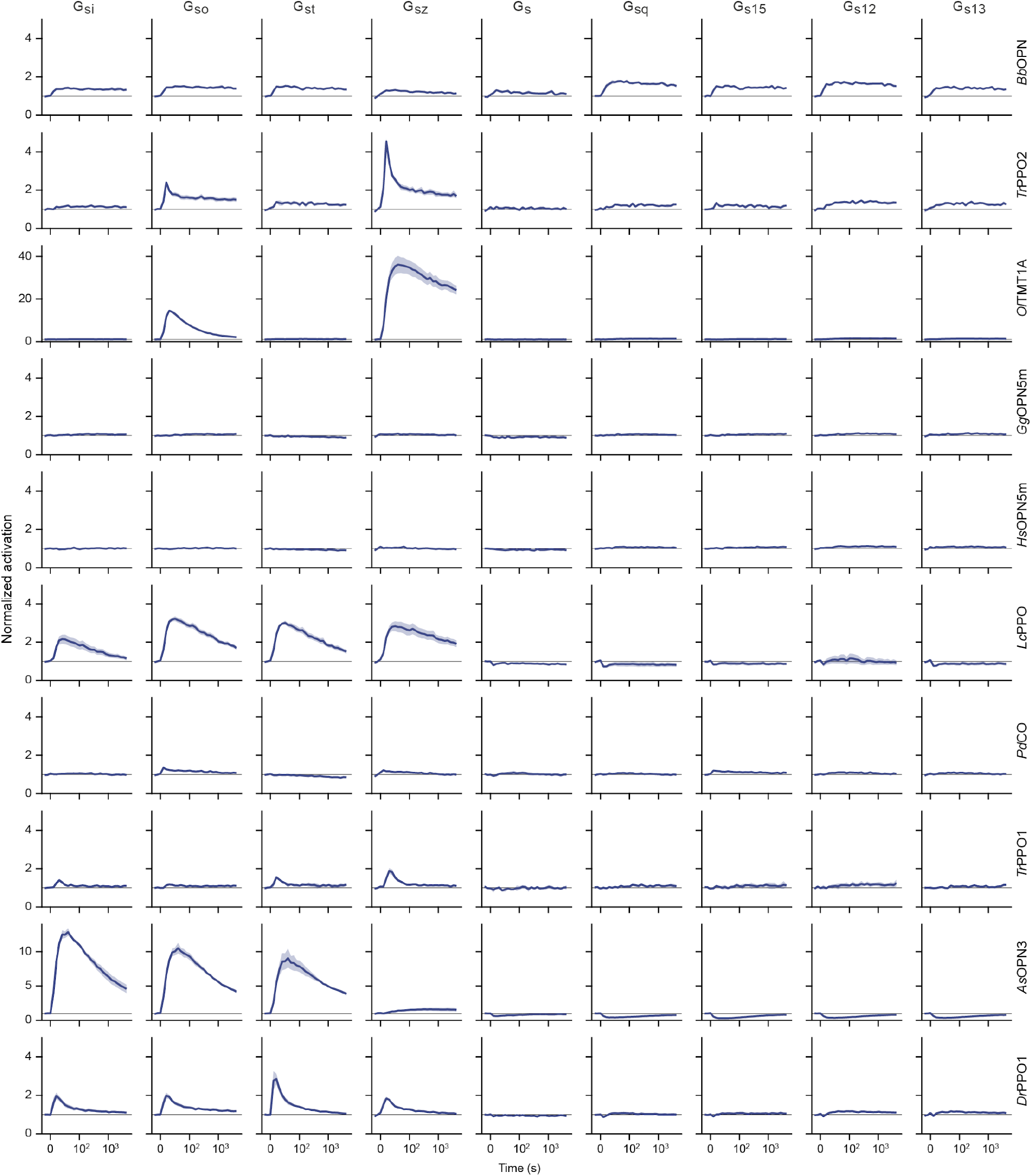
GsX assay of optoGPCR candidates. Time course of averaged bioluminescence reads for Gs-protein chimeras (columns) and optoGPCRs (rows). For *Ol*TMT1A, *Tr*PPO2, and *Bb*OPN a 1s 470nm light pulse (Time=0s) was used for optoGPCR activation, while all other optoGPCRs were activated with a 1s 365nm light pulse. Please note different scales for *As*OPN3 and *Ol*TMT1A. All data is shown as mean ± SEM (n= 2-6).

**Extended Data Fig. 3:**
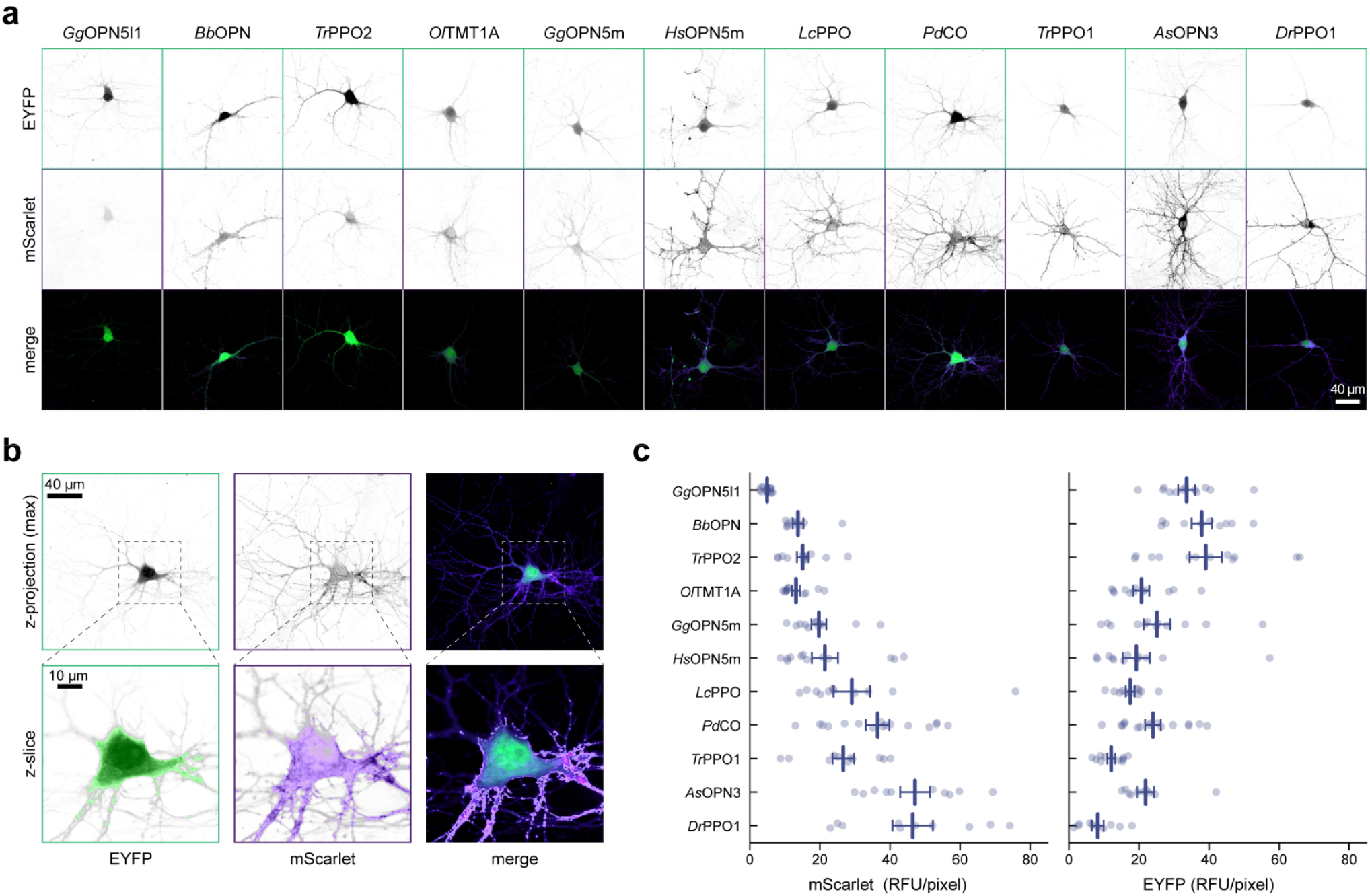
Quantification of optoGPCR expression in cultured neurons. **a**, Representative maximum intensity projection confocal images of neurons expressing the optoGPCRs together with a cytosolic EYFP. From top to bottom row: EYFP, mScarlet and merge of both signals. Inverted grayscale images are gamma corrected (1.25). The merged projections are false color coded by green (EYFP) and duo intense purple (mScarlet) lookup tables. **b**, *Pd*CO example for quantification of fluorescence measurements. Zoomed in single z-slices are shown for each channel in the lower row together with the calculated binary masks. To quantify the membrane expression only (bottom row, right), the calculated mask for the cytosolic EYFP channel was subtracted from the mScarlet channel mask. Color coding and lookup tables as in a, but gamma (1.75) for single z-slice images. **c**, Quantification of total mScarlet (left) and EYFP (right) fluorescence mean gray intensities, determined from equatorial z-slices. n=10-16. All data is shown as mean ± SEM.

**Extended Data Fig. 4:**
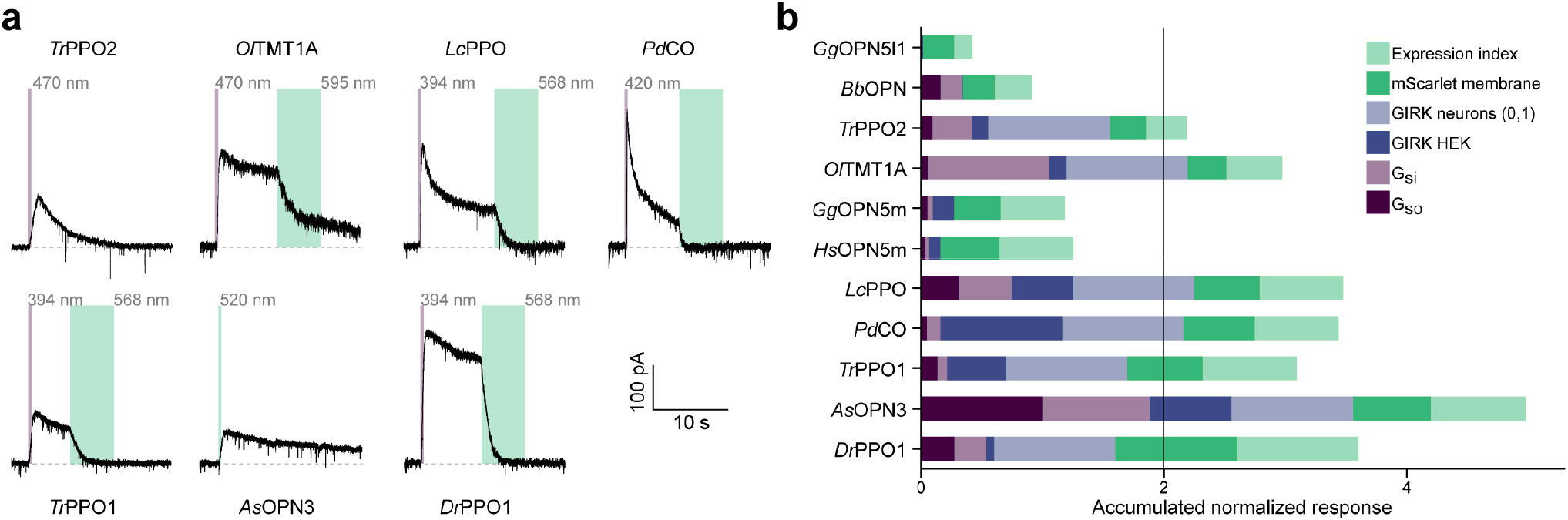
OptoGPCR GIRK coupling and benchmark summary. **a**, Representative whole-cell patch-clamp recordings of cultured neurons co-transfected with the different optoGPCRs and GIRK2.1, respectively. Neurons were kept at -70mV holding potential and a 0.5s light pulse at indicated wavelengths was used for optoGPCR activation, while 5s light pulse at indicated wavelengths was used for optoGPCR inactivation. Illumination intensities were adjusted to equal photon flux density for all wavelengths. No response to light could be recorded for *Gg*OPN5l1, *Bb*OPN, *Gg*OPN5m, and *Hs*OPN5m (data not shown). **b**, Accumulated normalized response across all assays used to benchmark optoGPCR candidates against each other. Within each assay, the mean value of each optoGPCR was normalized to the maximum mean value within the regarding assay. For positive GIRK-coupling shown in a, a value of 1 (0, in case no coupling detected) was assigned. Logarithmic GsX mean values were used. An arbitrary cutoff of 2 was chosen for the follow up benchmark in autaptic neurons.

**Extended Data Fig. 5:**
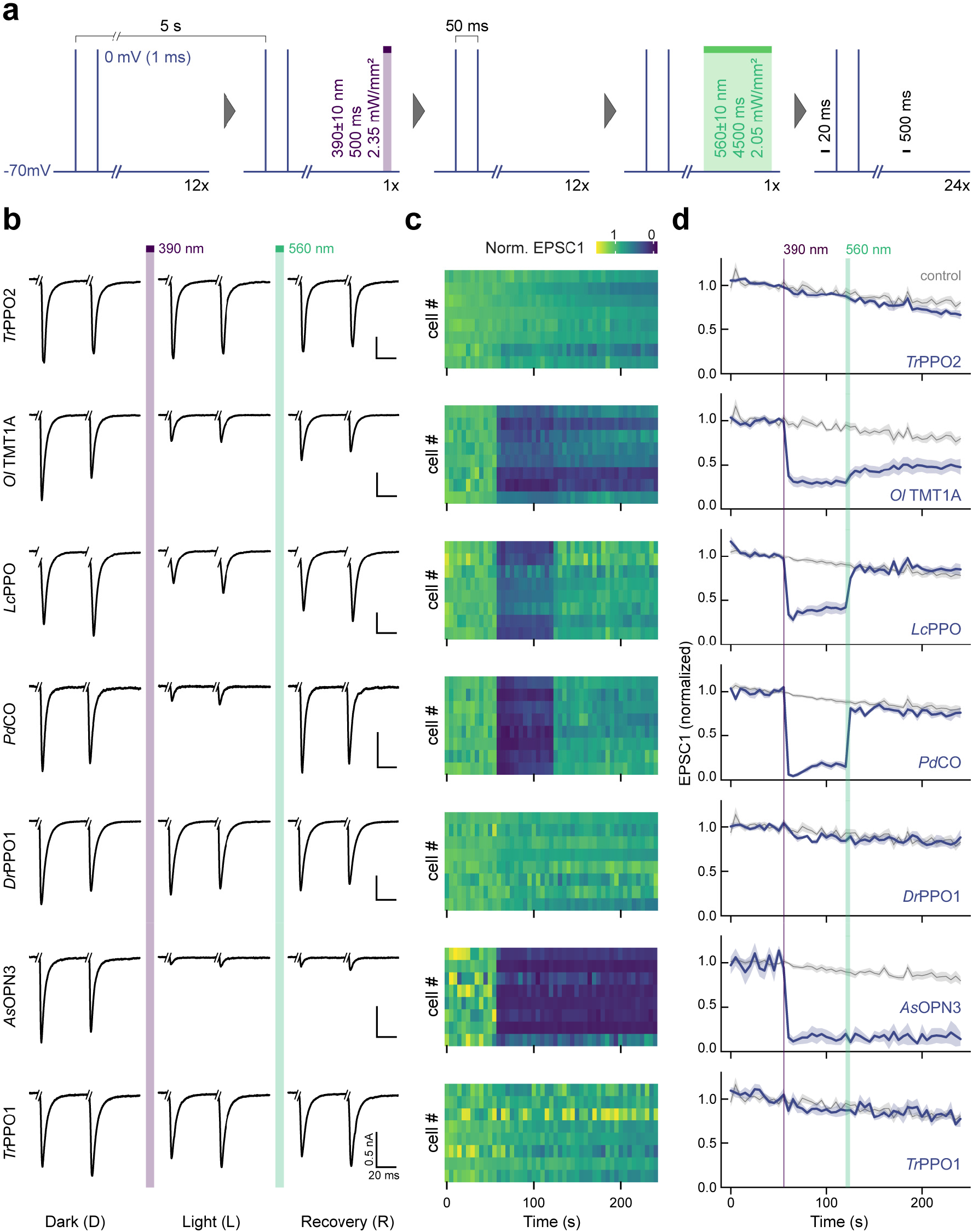
Inhibition of synaptic transmission in autaptic neurons. **a**, Experimental scheme of EPSC recordings in autaptic neurons. **b**, Representative averaged EPSC traces for the 7 investigated optoGPCR candidates recorded as described in a. Traces show 5 averaged sweeps. Current injections were cut for representation. **c**, Contour plot of EPSC amplitudes for 8 biological replicates. Amplitudes were normalized to the average of 5 EPSC1s prior 390 nm illumination. **d**, Timeplot of averaged EPSC data (across replicates) as shown in c, together with non-expressing control cells from matching autaptic cultures measured with the same protocol. Data shown as mean ± SEM (n=8).

**Extended Data Fig. 6:**
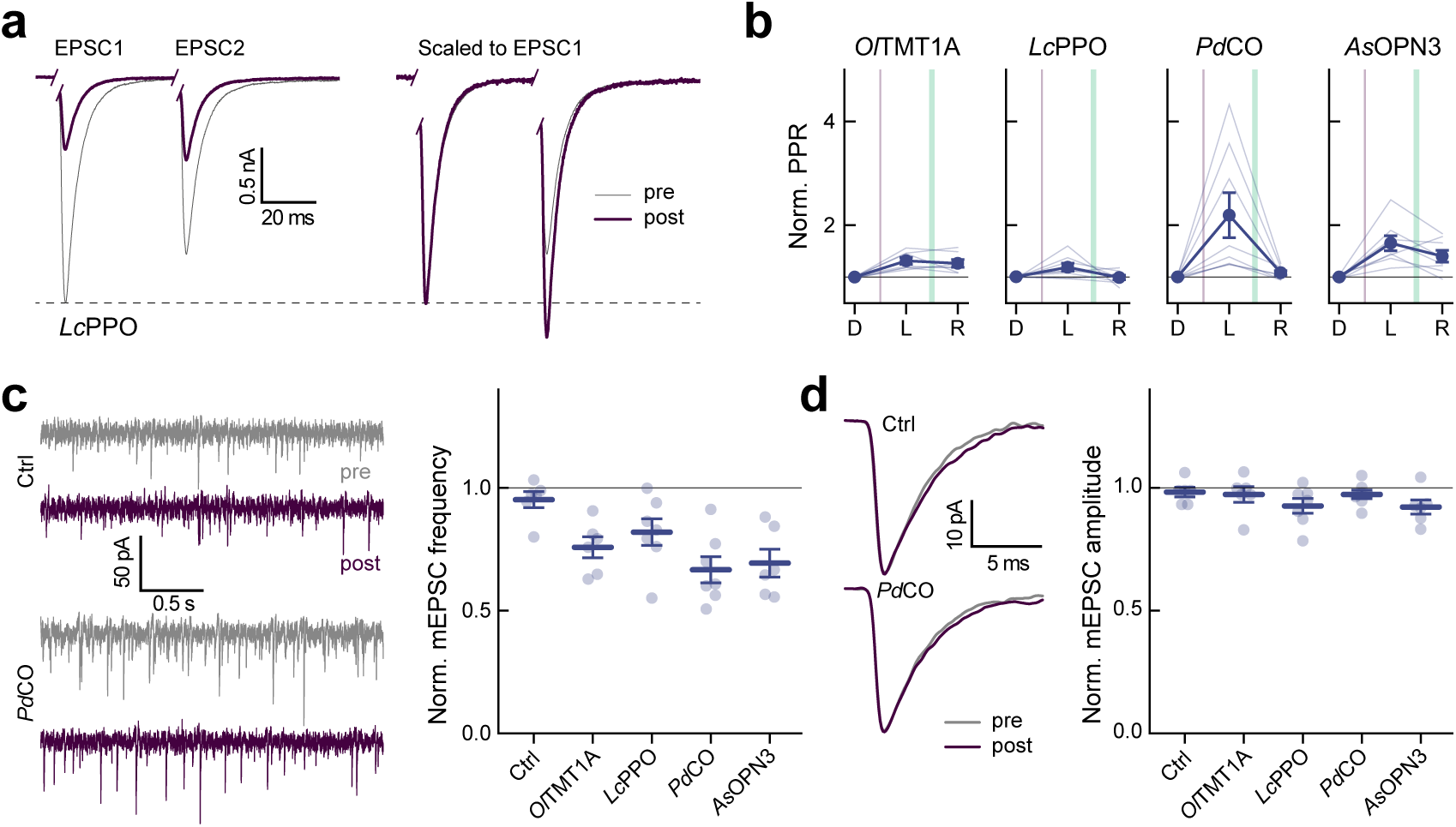
Paired-pulse recordings and mEPSCs in autaptic neurons. **a**, Paired-pulse recording of an *Lc*PPO expressing neuron (left), scaled to the first EPSC (right). **b**, Quantification of paired-pulse ratio (EPSC2/EPSC1). n= 8 **c**, Representative traces of mEPSCs (left) in a non-transduced control neuron (top) and a PdCO expressing autaptic cell (bottom). Right: Quantification of the mEPSC frequency after the light flash, normalized to the frequency prior light. n= 6-7. **d**, Quantal EPSC amplitude in control (top trace) and *Pd*CO-expressing (bottom trace) neurons pre and post illumination. Right: Quantification of the mEPSC amplitudes after the light flash, normalized to pre illumination; n= 6-7. All data is shown as mean ± SEM.

**Extended Data Fig. 7:**
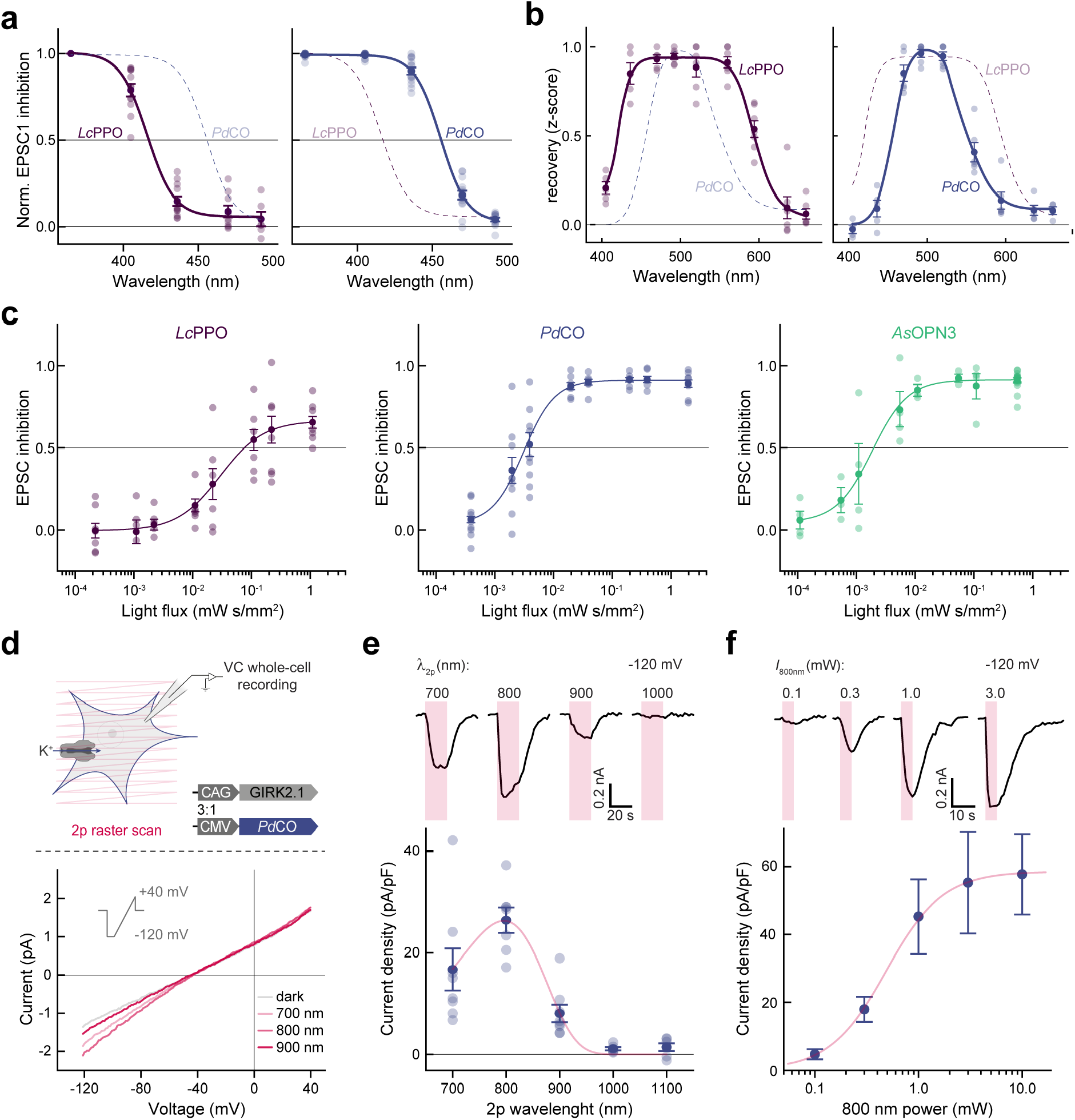
Biophysical properties in autaptic neurons. **a**, Quantification of EPSC inhibition versus applied wavelengths for *Lc*PPO (right) and *Pd*CO (left) including single cell measurement data points. For each cell, the inhibition was normalized to the maximum inhibition. Lines show dose-response fits. n= 6-15. **b**, Wavelength sensitivity of light induced recovery including single cell measurement data for *Lc*PPO (right) and *Pd*CO (left). n= 4-7. **c**, Quantification of absolute inhibition for the first EPSC post sample illumination versus light flux for (from left to right) *Lc*PPO (365 nm), *Pd*CO (405 nm) and *As*OPN3 (520 nm) including single cell measurement data points. Lines depict sigmoidal fits to calculate half maximal inhibition light flux (EC_50_) values. n= 3-17. **d**, Top: schematic of 2-photon (2p) recordings in HEK293T cells co-transfected with *Pd*CO-mScarlet and GIRK2.1 in a 1:3 ratio. Whole-cell patch-clamp recordings were performed and PdCO was activated with 2p raster scanning. Bottom: example recordings at various 2p wavelengths while applying a voltage ramp from -120 to +40mV. **e**, Top: example recordings of one representative HEK293T cell at -120mV holding potential for different scan wavelengths as indicated. Bottom: Quantification of 2p wavelength sensitivity; n= 6-8. Data was fitted with a 3-parametric Weibull distribution (red line) **f**, Top: example recordings of arepresentative HEK293T cell at -120mV for different 2p intensities of 800 nm. Bottom: Quantification of 2p sensitivity at 800 nm; n= 6-8. Data was fitted with a logistic dose response curve to determine the EC_50_ value (red line). All data is shown as mean ± SEM.

**Extended Data Fig. 8:**
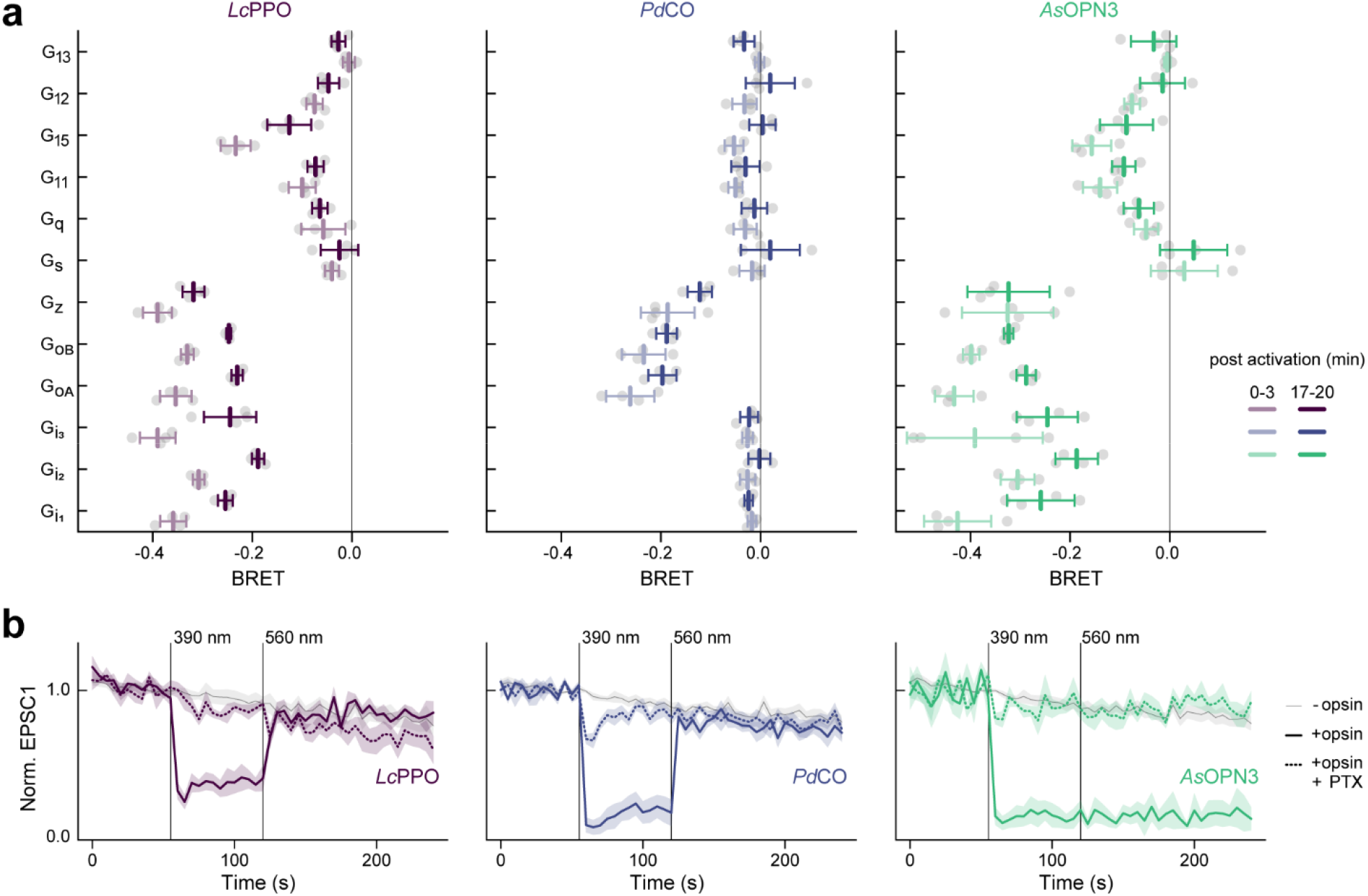
TRUPATH assay and pertussis toxin treatment in autaptic cultures. **a**, TRUPATH netBRET data for all G-protein classes integrated over 3 minutes post optoGPCR activation (light colors) and 17-20 minutes post activation (saturated colors). n= 4. **b**, EPSC recordings in autaptic neurons as described for initial benchmark of optoGPCRs with additional pertussis toxin (PTX) treatment. PTX strongly reduces optoGPCR-mediated EPSC inhibition, and therefore indicates a G_i/o_-dependent mechanism. n= 6-8. All data is shown as mean ± SEM.

**Extended Data Fig. 9:**
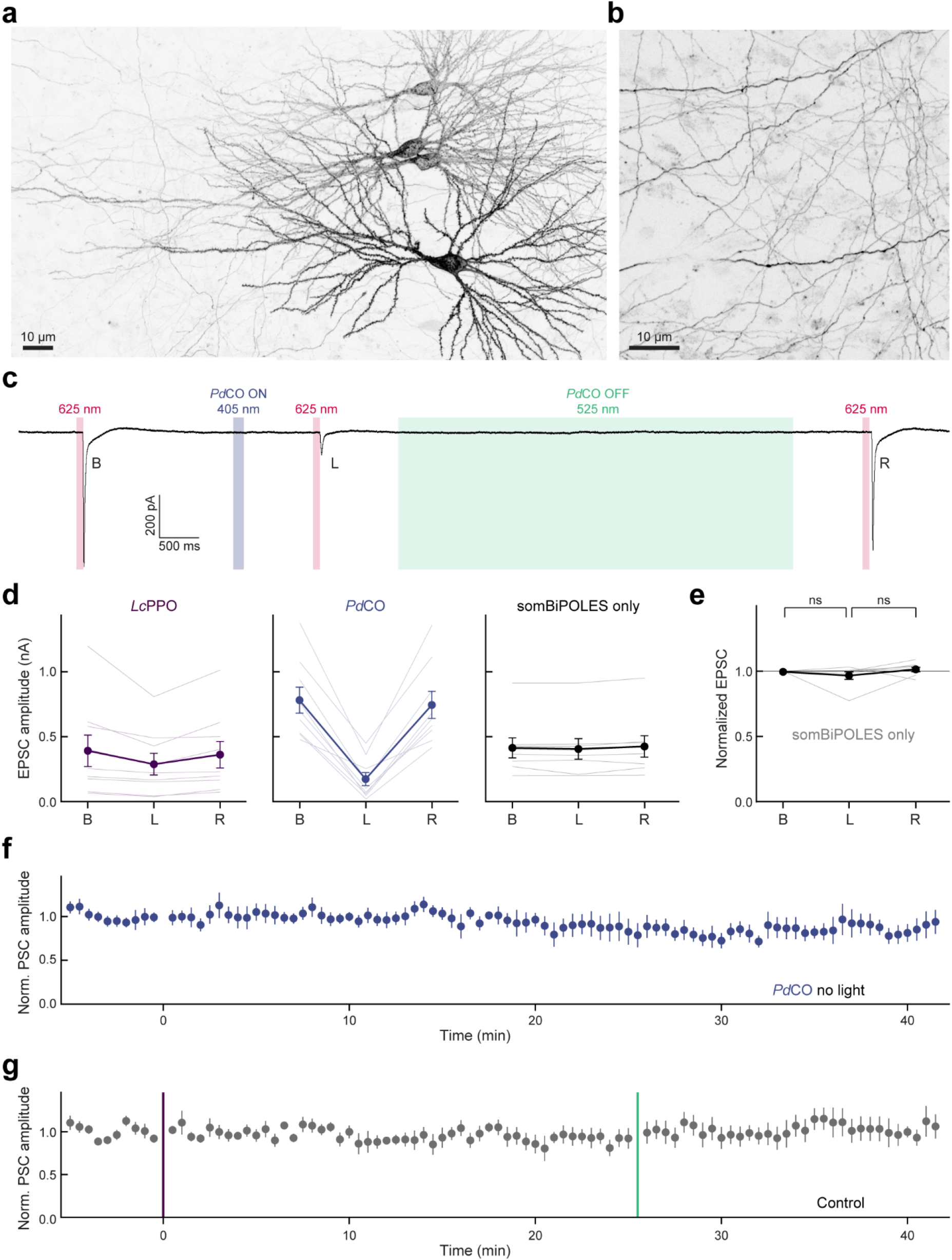
Electrophysiological measurements in organotypic slices. **a**, Two-photon maximum-intensity projections of CA3 neurons expressing *Pd*CO-mScarlet (delivered via electroporation). **b**, Axons (originating from CA3 neurons shown in a) projecting into stratum radiatum of CA1. **c**, Experimental design, and representative current trace for bidirectional optogenetic control of synaptic transmission at Schaffer collateral synapses using somBiPOLES and *Pd*CO. **d**, Quantification of absolute PSC amplitudes from bidirectional optogenetic experiments. n= 8-9. **e**, Normalized PSC amplitudes from control cultures expressing somBiPOLES alone, same as right panel in b. n= 8. **f**, Time course of normalized PSC amplitudes from control cultures expressing PdCO alone, without light stimulation. n= 4-6. **g**, Time course of normalized PSC amplitudes from control, non-expressing cultures, before and after light stimulation. n= 5-6. All data is shown as mean ± SEM.

**Extended Data Fig. 10:**
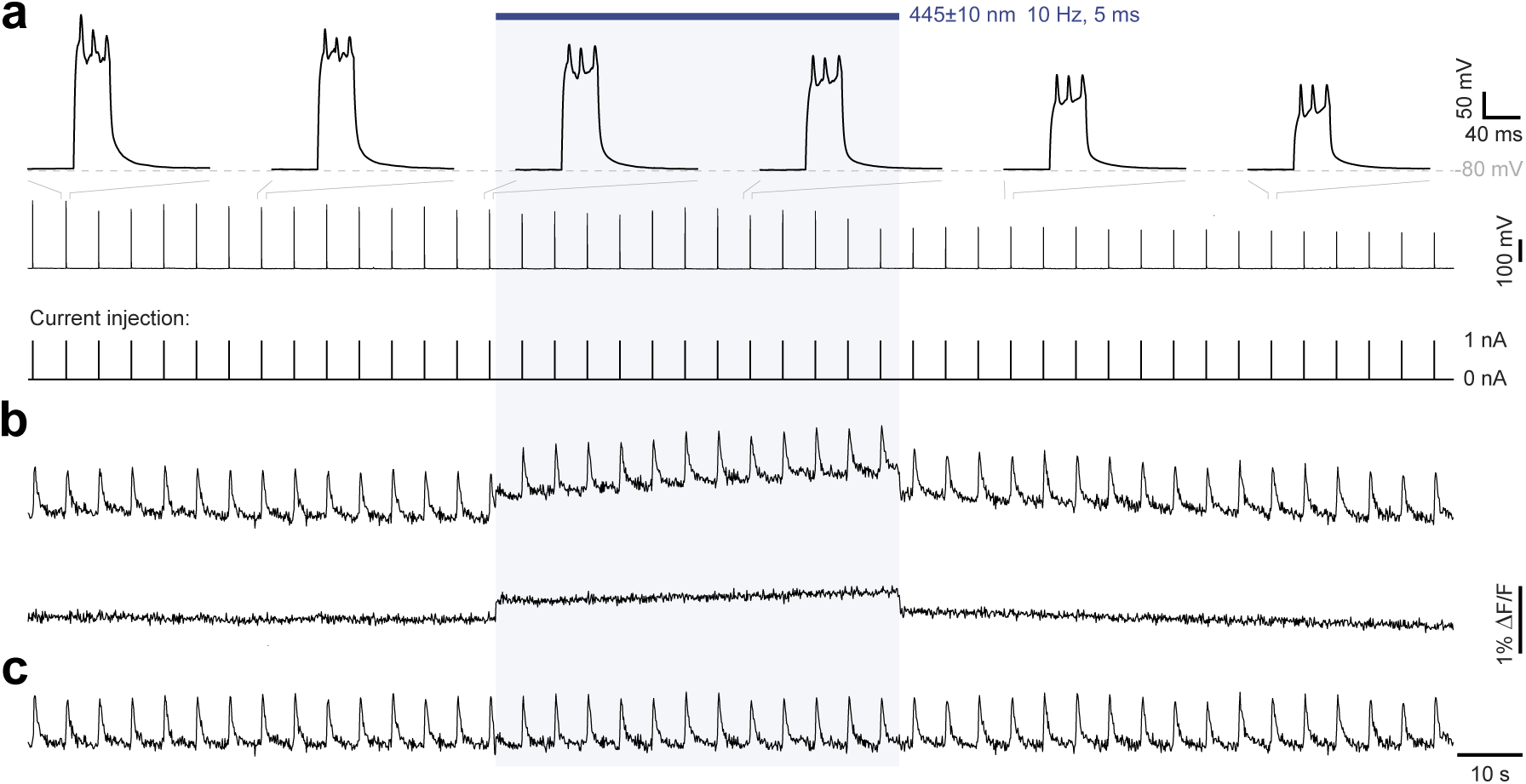
cell-attached calcium imaging and blue-light fluorescence increase. **a**, Voltage recording under tight-seal current-clamp conditions. 1 nA current pulses (40ms duration) were injected through the intact membrane patch at 0.2 Hz to evoke a train of action potentials. Top: magnified voltage recordings at selected time points. Middle: voltage recording of the complete experiment. Bottom: current injection. **b**, Top: raw ΔF/F signal from the neuron shown in a. Bottom: raw ΔF/F signal from a neuron in the same field of view recorded simultaneously. **c**, manually corrected ΔF/F signal as shown in b, top.

